# Dissection of insular cortex layer 5 reveals two sublayers with opposing modulatory roles in appetitive behavior

**DOI:** 10.1101/2022.06.15.494020

**Authors:** Makoto Takemoto, Shigeki Kato, Kazuto Kobayashi, Wen-Jie Song

## Abstract

The insular cortex (insula) is known to play a modulatory role in motivated behaviors including feeding and drinking. Previous studies have revealed that the anterior and posterior subregions of the insula have differential subcortical efferents and roles, yet the anatomical and functional heterogeneity among the cortical layers remains poorly understood. Here, we show that layer 5 of the mouse dysgranular insula has two distinct neuronal subpopulations along the entire anterior-posterior axis: the upper layer (L5a) population, expressing NECAB1, projects bilaterally to the lateral and capsular divisions of the central amygdala, and the deeper layer (L5b) population, expressing CTIP2, projects ipsilaterally to the parasubthalamic nucleus and the medial division of the central amygdala. Optogenetically activating L5a and L5b neuronal populations in thirsty mice led to suppressed and facilitated water spout licking, respectively, in a single-spout test. However, the opto-activation induced no avoidance against or preference for the spout paired with the opto-stimulation in a two-spout choice test, indicating no induction of emotional valences by the opto-activation per se. Our results suggest sublayer-specific bidirectional modulatory roles of insula layer 5 in the motivational aspect of appetitive behavior.

## Introduction

The insular cortex (insula) mediates top-down modulation of feeding and drinking behaviors in response to taste quality (1, 2), homeostatic/visceral states (3, 4), innate threat (3), and learned stimuli (5–10), as well as the modulation of emotional (11) and social (3, 12, 13) behaviors. In particular, the modulation of feeding or drinking is differentially regulated by distinct subregions of the insula along the anterior-posterior (A-P) axis (1), and region-specific projections to the amygdala complex are suggested to underlie the behavioral control. The projection of the anterior insula to the basolateral amygdala (BLA) and the projection of the posterior insula to the central amygdala (CeA) serve to enhance and suppress drinking, respectively, by inducing opposite emotional valences (2). The suppressive effect of the posterior insula-CeA circuit on drinking and feeding behaviors has been consistently observed in several studies (3, 8, 14). Because this region-specific feeding control suits well with topographic representations of taste qualities in the gustatory insula, where sweet and bitter (i.e., appetitive and aversive) tastes are primarily represented in the anterior and posterior part (15), respectively, the region-specific hard-wired connections of the insula are thought to underlie the taste quality-dependent control of feeding as an innate behavioral response.

Feeding, however, is not merely determined by taste quality but is also adaptively regulated by internal and external circumstances (16, 17). Therefore, additional neural substrates may also participate in the modulation of feeding. Indeed, Stern et al. (10) have demonstrated that the CeA projection from nitric oxide synthase-1-expressing neuronal subpopulation mediates an enhanced feeding in response to a learned context. This observation contrasts the feeding-suppressive effect of the insula-CeA circuit reported in the studies mentioned above, suggesting heterogeneity in neuronal populations of the posterior insula that project to the CeA.

Anatomical and functional heterogeneity of subcortical projection neurons of the insula has been investigated intensely in terms of the differences among the subregions (1–3, 13, 18–23). However, much less attention has been given to the layer specificity that includes neuronal diversity within a layer and among layers, regardless of the unique laminar cytoarchitecture of the insula (19, 24). Only a few anatomical studies have been carried out so far, reporting the layer 5 (L5) origin of insula projections to subcortical structures, such as the parabrachial nucleus of the pons (25) and the nucleus of the solitary tract (20, 26–28).

Here, we show that L5 of the dysgranular insula (DgI) has two distinct neuronal subpopulations along the whole A-P axis. One, primarily located in the upper L5 (L5a), projects bilaterally to the lateral and capsular divisions of the CeA (CeL/CeC) and suppresses appetitive drinking upon its optogenetic activation. The other, predominantly located in the deeper L5 (L5b), projects ipsilaterally to the parasubthalamic nucleus (PSTh) of the hypothalamus and the medial division of the CeA (CeM) and facilitates appetitive drinking upon its optogenetic activation. These findings suggest opposite modulatory roles of sublayers of insula L5 in appetitive behavior.

## Results

### Hemispheric difference in insula projections to the extended amygdala and parasubthalamic nucleus

To investigate the heterogeneity of the mouse insula subcortical projections, we first labeled neurons in the middle-posterior insula with GFP using the AAV2.CAG.GFP vector. The injection site (0.05 ± 0.3 mm posterior to Bregma) with GFP-expressing neuronal somata covered the ventral part of the granular insula (GI), which contains a clear granular layer (i.e., layer 4), and the DgI, with a vague layer 4, located dorsal to the agranular insula (AgI) that lacks a six-layered structure (19, 24) (Fig. 1A). GFP-labeled axon arbors were found in many subcortical structures that include the paraventricular and mediodorsal thalamic nuclei, nucleus reuniens, CeA, basomedial amygdala (BMA) (Fig. 1B), bed nucleus of the stria terminalis (BNST), interstitial nucleus of the posterior limb of the anterior commissure (Fig. 1E), parvicellular part of the ventral posterior thalamic nucleus, and PSTh (Fig. 1F), as well as the ventral striatum and brainstem, in line with the results of previous anatomical studies (2, 10, 18, 19, 23, 29). There were virtually no retrogradely labeled cell bodies in these structures.

**Figure 1.**
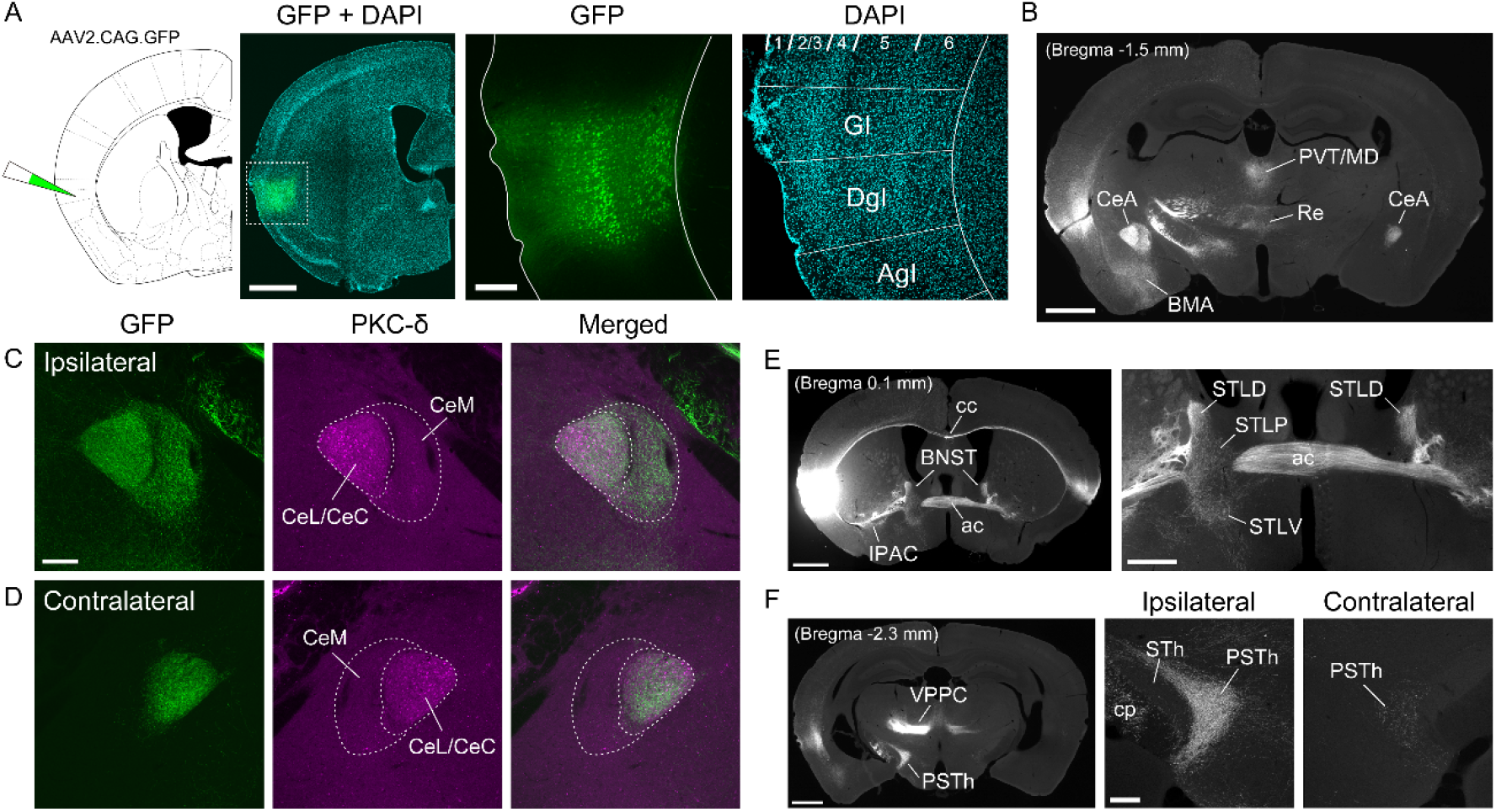
Hemispheric difference in subcortical projections of the insula. (A) A typical example of the insula injected with the AAV2.CAG.GFP vector. GFP is expressed primarily in the middle to deep layers of the dysgranular (DgI) and granular (GI) regions of the insula. (B) A gray-scale image of the distribution of GFP-expressing axon arbors in the amygdala and thalamus. (C) GFP-labeled axon arbors in the ipsilateral CeA. The arborization (green) is located in the PKC-positive CeL/CeC (magenta) and the adjacent CeM. (D) GFP-labeled axon arbors in the contralateral CeA. The arborization is localized in the PKC-δ positive CeL/CeC. (E) A gray-scale image of GFP-expressing axons in the BNST and ventral striatum (left) and a magnified view of the BNST (right). (F) A gray-scale image of GFP-expressing axon arbors in the PSTh and thalamus (left) and magnified views of the ipsilateral and contralateral PSTh (right). Abbreviations: ac, anterior commissure; AgI, agranular insula; BMA, basomedial amygdala; cc, corpus callosum; cp, cerebral peduncle; IPAC, interstitial nucleus of the posterior limb of the anterior commissure; MD, mediodorsal thalamic nucleus; PVT, paraventricular thalamic nucleus; Re, nucleus reuniens; STh, subthalamic nucleus; VPPC, parvicellular part of the ventral posterior thalamic nucleus. Scale bars: 1 mm (A-left, B and E-left, F-left), 0.2 mm (A-right, C and F-right), 0.5 mm (E-right).

Focusing on the dense labeling of GFP-positive axons projecting to the CeA of both hemispheres, we found the projection patterns to be different between the hemispheres. The identification of the CeL/CeC with a molecular marker, the protein kinase C delta (PKC-δ) (30), revealed that the ipsilateral projection was distributed in both the CeL/CeC and adjacent CeM (Fig. 1C), whereas the contralateral projection was confined to the CeL/CeC (Fig. 1D). Such asymmetric projections were also found in the lateral division of the BNST (STL), in which the ipsilateral projections were seen in the dorsal (STLD, also referred to as the oval nucleus of the BNST), posterior (STLP), and ventral (STLV) part of the STL, whereas the contralateral projection was found only in the STLD (Fig. 1E). In the hypothalamus, dense axon arbors expressing GFP were packed in the ipsilateral PSTh, but few labeled fibers were found on the contralateral side (Fig. 1F). These findings indicate that unilateral and bilateral subcortical projections of the insula are organized in a target-dependent fashion.

### Sublaminar and regional distribution of anatomically distinct neuronal populations in DgI L5

We hypothesized that the axons with ipsilateral projections and those with contralateral projections originate from distinct neuronal populations of the insula. To test this hypothesis, we performed dual injections of retrograde tracers cholera-toxin B subunit (CTB)-green and CTB-red into the right CeA–always covering at least a part of the CeL/CeC–and into the left posterior lateral hypothalamic area that includes the PSTh, respectively (Fig. 2A, Fig. S1A). We found CTB-green- and CTB-red-labeled cells in the DgI in both hemispheres with distinct laminar distributions in the left hemisphere (Fig. 2B, also see Fig. S1B vs. Fig. S1C). In the left insula, cells retrogradely labeled from the contralateral (i.e., right) CeA (CeA-contra) were distributed primarily in L5a and sparsely in L5b (Fig. 2C, D, green), whereas cells retrogradely labeled from the ipsilateral (i.e., left) PSTh (PSTh-ipsi) were predominantly located in L5b (Fig. 2C, D, red) (also see Fig. S1D). We found no double-labeled cells: 0 double-labeled cells out of 756 and 1016 cells retrogradely labeled by CTB injections into the CeA-contra and PSTh-ipsi, respectively, analyzed in 10 coronal sections from 5 brains. This suggests that the contralateral CeL/CeC and ipsilateral PSTh (Fig. 1) receive input from non-overlapping neuronal subpopulations in L5 of the DgI. To further characterize the sublaminar distribution of the subpopulations, we investigated the expression of molecular markers of L5 sublayers, namely, the N-terminal EF-hand calcium-binding protein 1 (NECAB1) for pyramidal neurons in L5a (31) and the chicken ovalbumin upstream promoter transcription factor-interacting proteins 2 (CTIP2) for pyramidal tract-type neurons in L5b (32, 33) (Fig. S1E, F). Immunohistochemical analyses showed that the majority of CeA-contra-projecting (87.1% on average, n = 3 mice) but virtually no PSTh-ipsi-projecting (0.7% on average, n = 3 mice) CTB-labeled cells were positive for NECAB1 (Fig. 2E, G), while almost all PSTh-ipsi-projecting (98.1% on average, n = 3 mice) but almost no CeA-contra-projecting (0.3 % on average, n = 3 mice) CTB-labeled cells were positive for CTIP2 (Fig. 2F, G). These results indicate that neurons contralaterally projecting to the CeL/CeC and those ipsilaterally projecting to the PSTh have distinct L5 sublayer-specific molecular profiles.

**Figure 2.**
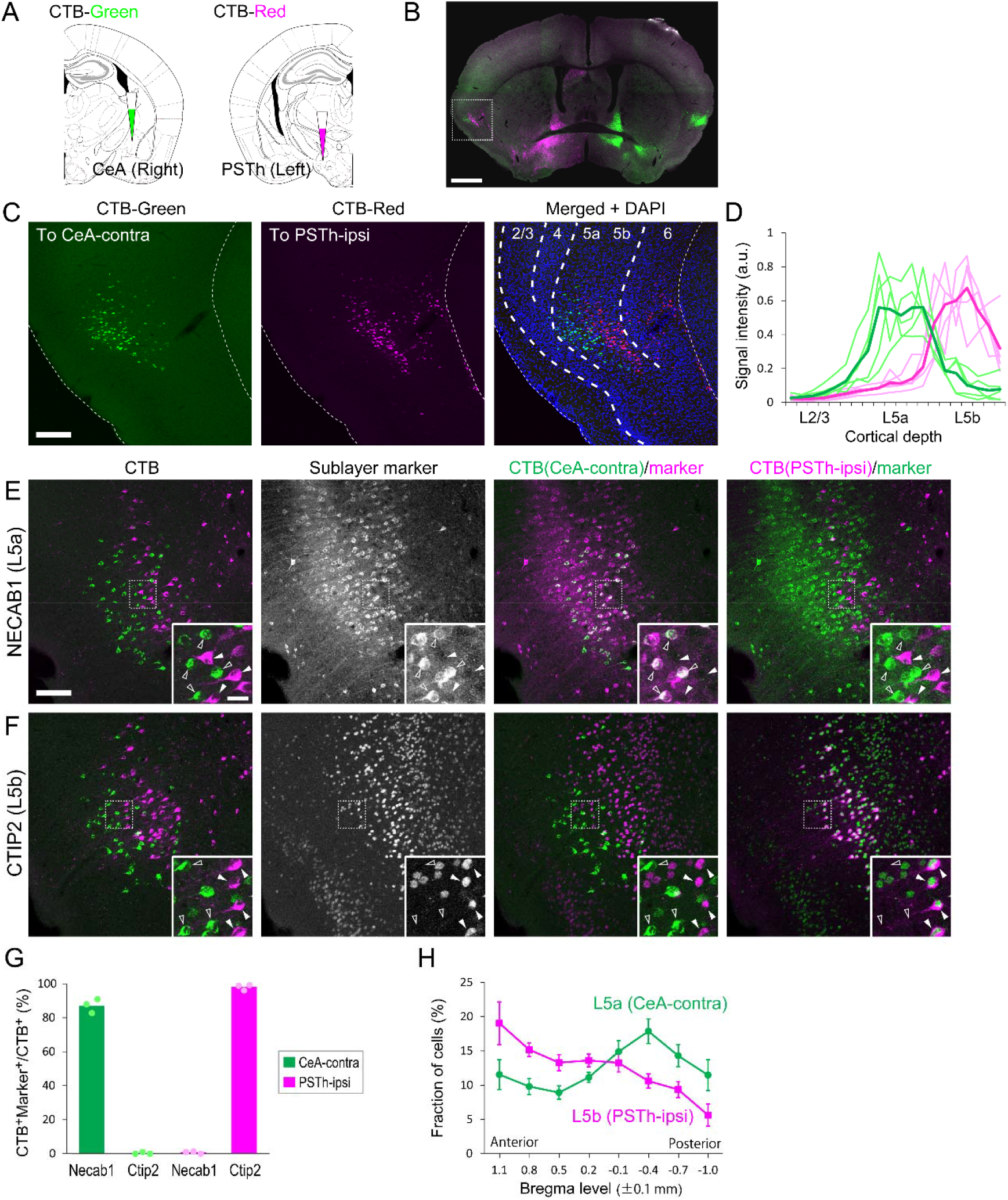
Sublaminar and regional distribution of anatomically distinct cell populations in L5 of the insula. (A) Illustrations depicting dual-color CTB injections into the right CeA and left PSTh. (B) A typical example of the confocal image of a coronal section containing CTB-labeled cells in the insula. (C) Higher magnification views of the left insula (boxed region in B). (D) Relative fluorescence intensities (arbitrary unit) of CTB labeling across cortical depth in the DgI (L2/3 to L5b). Note that the density of labeled cells across cortical depth can be estimated by the averaged fluorescence because of the localized fluorescence to cell somas. The signal intensities of CTB-green-labeled cells (projecting to the CeA-contra) and CTB-red-labeled cells (projecting to the PSTh-ipsi) are shown in green and magenta, respectively. Light colors represent individual mice (n = 5 mice with dual CTB injections), and dark colors are the means. (E) Fluorescent images of a section containing CTB-labeled cells with immunostaining of NECAB1 (an L5a marker, gray-scale). CeA-contra-projecting CTB-labeled cells (open arrowheads) but not PSTh-ipsi-projecting CTB-labeled cells (solid arrowheads) are immunopositive for NECAB1. (F) Fluorescent images of a section containing CTB-labeled cells with immunostaining of CTIP2 (an L5b marker, gray-scale). PSTh-ipsi-projecting CTB-labeled cells (solid arrowheads) but not CeA-contra-projecting CTB-labeled cells (open arrowheads) are immunopositive for CTIP2. (G) The proportion of anatomically distinct CTB-labeled cell populations positive for the sublayer-specific molecular markers. Dots represent individual mice. Bars indicate the means (87.1% for NECAB1/CeA-contra; 0.33% for CTIP2/CeA-contra; 0.75% for NECAB1/PSTh-ipsi; 98.1% for CTIP2/ PSTh-ipsi, n = 3 mice). (H) The fraction of CeA-contra-projecting CTB-labeled cells (L5a population, green) and PSTh-ipsi-projecting CTB-labeled cells (L5b population, magenta) at eight locations across the A-P axis of the DgI. The A-P level of the maximum proportion is 0.4 ± 0.1 mm posterior to Bregma for the L5a population (one-way ANOVA, F(7,40) = 20.2, p = 2.75 × 10^−11^; Dunnett’s post hoc test, p < 0.05 [vs. −0.1 ± 0.1 mm]; p < 0.01 [vs. −0.7 ± 0.1 mm]; p < 0.001 [vs. 1.1, 0.8, 0.5, 0.2, −1.0 ± 0.1 mm], n = 6 mice) and 1.1 ± 0.1 mm anterior to Bregma for the L5b population (one-way ANOVA, F(7,32) = 32.9, p = 7.22×10-13; Dunnett’s post hoc test, p < 0.01 [vs. 0.8 ± 0.1 mm]; p < 0.001 [vs. 0.5, 0.2, −0.1, −0.4, −0.7, −1.0 ± 0.1 mm], n = 5 mice). Error bars represent standard deviations. Scale bars: 1 mm (B), 0.2 mm (C), 0.1 mm (E), 20 μm (inset in E).

As the insula exhibits region specificity in subcortical projections (2, 3, 23), we examined the distribution of L5 subpopulations along the A-P axis. We found CeA-contra-projecting cells (the L5a population) in the entire DgI with the peak proportion of the cells at the middle of the posterior insula (0.4 ± 0.1 mm posterior to Bregma) (Fig. 2H, green). PSTh-ipsi-projecting cells (the L5b population) were found in the entire DgI as well; however, the proportion of the cells was highest at the most anterior part we analyzed (1.1 ± 0.1 mm anterior to Bregma), which gradually decreased posteriorly (Fig. 2H, red), suggesting an A-P gradient in the DgI. Both cell populations thus existed across the entire DgI with distinct A-P distributions.

Unlike the retrograde labeling in the contralateral CeA, retrograde labeling in the ipsilateral CeA resulted in cells distributed more broadly in the DgI layers (Fig. S1C). Specifically, when the ipsilateral injection covered primarily the CeL/CeC, most retrogradely labeled cells were found in L5a and L2/3, and some of the L5a cells were double-labeled by the CTB injection into the contralateral CeA (Fig. S2A, B). When the ipsilateral CTB injection mainly covered the CeM, most retrogradely labeled cells were found in L5b and L2/3, and only few cells were labeled in L5a (Fig. S2A, C). These results suggest a layer-specific ipsilateral DgI input to the CeA: the CeL/CeC receives input from L5a and L2/3, while the CeM receives input from L5b and L2/3.

To reveal the projection targets for axon collaterals of the CeA-contra-projecting L5a population and PSTh-ipsi-projecting L5b population, we expressed GFP selectively in each subpopulation and examined the distribution of the labeled axon arborizations. For GFP labeling of the L5a population, we combined the injection of a retrograde AAV carrying the Cre recombinase gene (AAVretro.EF1a.mCherry-ires-Cre) (34) into the right CeA with the injection of an AAV encoding Cre-inducible GFP (AAV2.CAG.FLEX.GFP) into the left insula for GFP expression in the Cre-expressing neurons (Fig. 3A, B). We observed GFP-expressing cell somas primarily in L5a of the left DgI, most (but not all) of which were also detectable for mCherry (the Cre expression indicator) (Fig. 3B), and GFP-expressing axon arbors in the CeL/CeC in the hemisphere contralateral to the GFP-expressing insula (Fig. 3C). Like the contralateral projection, the ipsilateral GFP-labeled axon arbors in the amygdala were also confined to the CeL/CeC (Fig. 3D, E; Fig. S3A), indicating bilateral projections of the L5a subpopulation to the CeL/CeC. Moreover, we also found bilateral projections of GFP-labeled axons in the BNST, where the axon arbors were restricted to the STLD and juxtacapsular part of the BSTL (STLJ) (Fig. S3C).

**Figure 3.**
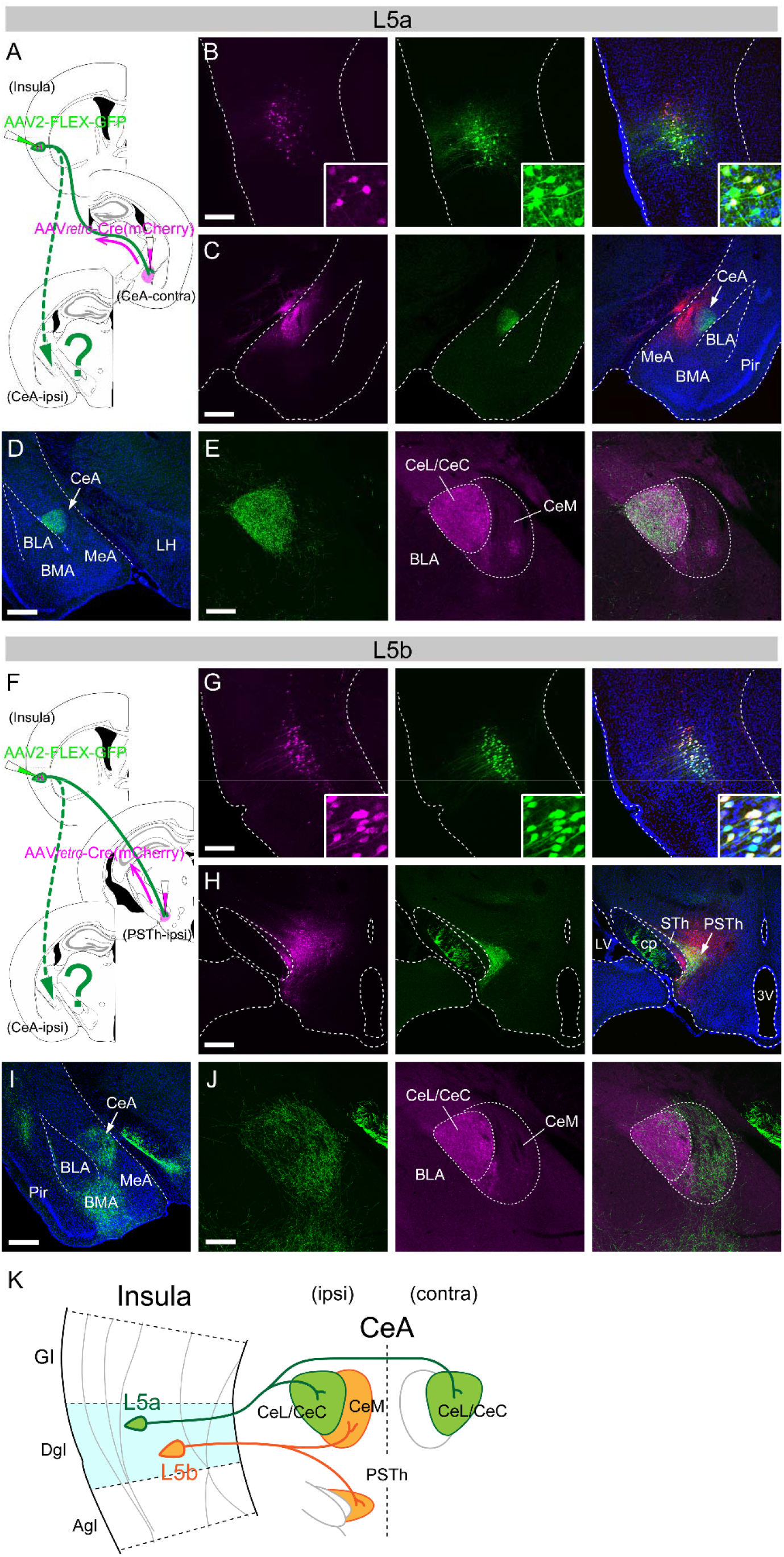
Differential organizations of axonal projections of DgI-L5 subpopulations to the CeA. (A) Illustrations depicting AAV injections for Cre-induced GFP expression in the L5a population. Note that a retrograde AAV carrying Cre was injected into the CeA contralateral to the insula injected with an AAV carrying Cre-inducible GFP. (B) Fluorescent images of the AAV-injected site in the insula. Left: mCherry (a Cre expression reporter), middle: GFP, right: merged (also counterstained with DAPI, blue). (C) Fluorescent images of the AAV-injected site in the CeA. Left: mCherry (indicative of the AAV injection), middle: GFP (axon arbors restricted to the CeL/CeC), right: merged (also with DAPI). (D) GFP-expressing axon arbors in the amygdala ipsilateral to the insula injected with the AAV vector. (E) A higher magnification view of the CeA ipsilateral to the insula with the AAV injection. Left: GFP (axon arbors restricted to the CeL/CeC), middle: anti-RGS14 immunostaining identifying the CeL/CeC but not CeM, right: merged. (F) Illustrations depicting AAV injections for Cre-induced GFP expression in the L5b population. (G) Fluorescent images of the AAV-injected site in the insula. Left: mCherry (Cre), middle: GFP, right: merged (also with DAPI). (H) Fluorescent images of the AAV-injected site in the ipsilateral hypothalamus. Left: mCherry (AAV injected site), middle: GFP (axon arbors restricted to the PSTh), right: merged (also with DAPI). (I) GFP-expressing axon arbors in the amygdala ipsilateral to the insula with the AAV injection. Labeled axons are found in the CeA and BMA. (J) A higher magnification view of the ipsilateral CeA. Left: GFP (axon arbors predominantly located in the CeM), middle: anti-RGS14 immunostaining, right: merged. Scale bars: 0.2 mm (B, F, and D-right, H-right), 0.5 mm (C, G, and D-left, H-left). (K) Schematic diagram of the summary for CeA projections of DgI-L5 neuronal subpopulations. Abbreviations: 3V, third ventricle; BLA, basolateral amygdala; BMA, basomedial amygdala; cp, cerebral peduncle; LH, lateral hypothalamus; LV, lateral ventricle; MeA, medial amygdala; Pir, piriform cortex; STh, subthalamic nucleus.

We took a similar strategy to label the axon collaterals of the L5b population. Specifically, we made injections of the AAVretro.EF1a.mCherry-ires-Cre into the PSTh and the AAV2.CAG.FLEX.GFP into the insula of the same hemisphere (Fig. 3F). Besides GFP-expressing cell somas in L5b of the insula, we found, in the ipsilateral hemisphere, dense axon arbors in the PSTh (Fig. 3G, H) and CeM and BMA of the amygdala, but only sparse terminals in the CeL/CeC (Fig. 3I, J; Fig. S3B). Thus, the organizations of ipsilateral projections originating from L5a and L5b subpopulations were largely complementary within the CeA, while the contralateral projection originated only from the L5a population and was confined to the CeL/CeC (Fig. 3K). These results are compatible with our observation that the ipsilateral CTB injection biased to the CeM preferentially labeled the L5b cells, as well as L2/3 cells (Fig. S2C), while the injection biased to the CeL/CeC preferentially labeled the L5a cells, as well as L2/3 cells (Fig. S2B) in the DgI. In the BNST, we found that GFP-labeled axon arbors originating from the L5b population were localized in the STLP and STLV but not in the STLD/STLJ, only in the ipsilateral hemisphere (Fig. S3D).

### Opposing modulatory roles of DgI L5 sublayers on appetitive behavior

The anatomical findings described above strongly suggest a functional difference between the two subpopulations of L5 neurons in the DgI. Recent studies have reported that the insula bidirectionally modulates drinking behavior in a region-specific manner (1, 2), while the CeL bidirectionally regulates feeding in a neuronal subtype-specific fashion (35–38). Additionally, the CeC and CeM were reported to play opposing roles in appetitive behaviors (35), and the PSTh was shown to participate in the processing of palatability (29) or, on the contrary, the suppression of feeding (38–42). These findings prompted us to ask whether the sublayers of the DgI L5 play distinct roles in the modulation of appetitive behavior. Therefore, we examined the effect of optogenetic activation of each sublaminar neuronal population on drinking behavior in thirsty mice, a model of appetitive behavior. We expressed Channelrhodopsin-2 (ChR2) (43) fused with tdTomato in either the L5a population (bilaterally projecting to the CeL/CeC) or the L5b population (ipsilaterally projecting to the PSTh and CeM) using a retrograde AAV (AAVretro.CAG.hChR2-H134R-tdTomato). We made a unilateral injection of the virus: the contralateral CeA injection for the L5a population activation or ipsilateral PSTh injection for the L5b population activation (Fig. 4A), and illuminated only one hemisphere with sublayer-specific ChR2 expression, because bilateral injections of the ChR2 virus into the CeA can cause ChR2 expression in multiple DgI layers (i.e., L2/3, L5a, and L5b) that contain both ipsilaterally and contralaterally projecting neuronal populations (Fig. S2) and make it practically impossible to optogenetically activate the L5a or L5b populations selectively. To optically stimulate ChR2-expressing neurons, a chip light-emitted diode (LED) was embedded over a small cranial window above the middle-posterior insula and was wirelessly controlled to deliver blue light to the cortex with the dura mater intact (Fig. 4A). This method was thus noninvasive to the cortex. The effectiveness of optogenetic stimulation was confirmed by detecting c-Fos expression in the DgI, induced by repetitive LED illuminations at the end of all behavioral experiments in each mouse: c-Fos immunoreactivity was primarily found in the ChR2-expressing layer of the stimulated side of the DgI (Fig. S4A, C), and the number of c-Fos positive cells in the stimulated side was significantly higher than in the non-stimulated side (Fig. S4B, D).

**Figure 4.**
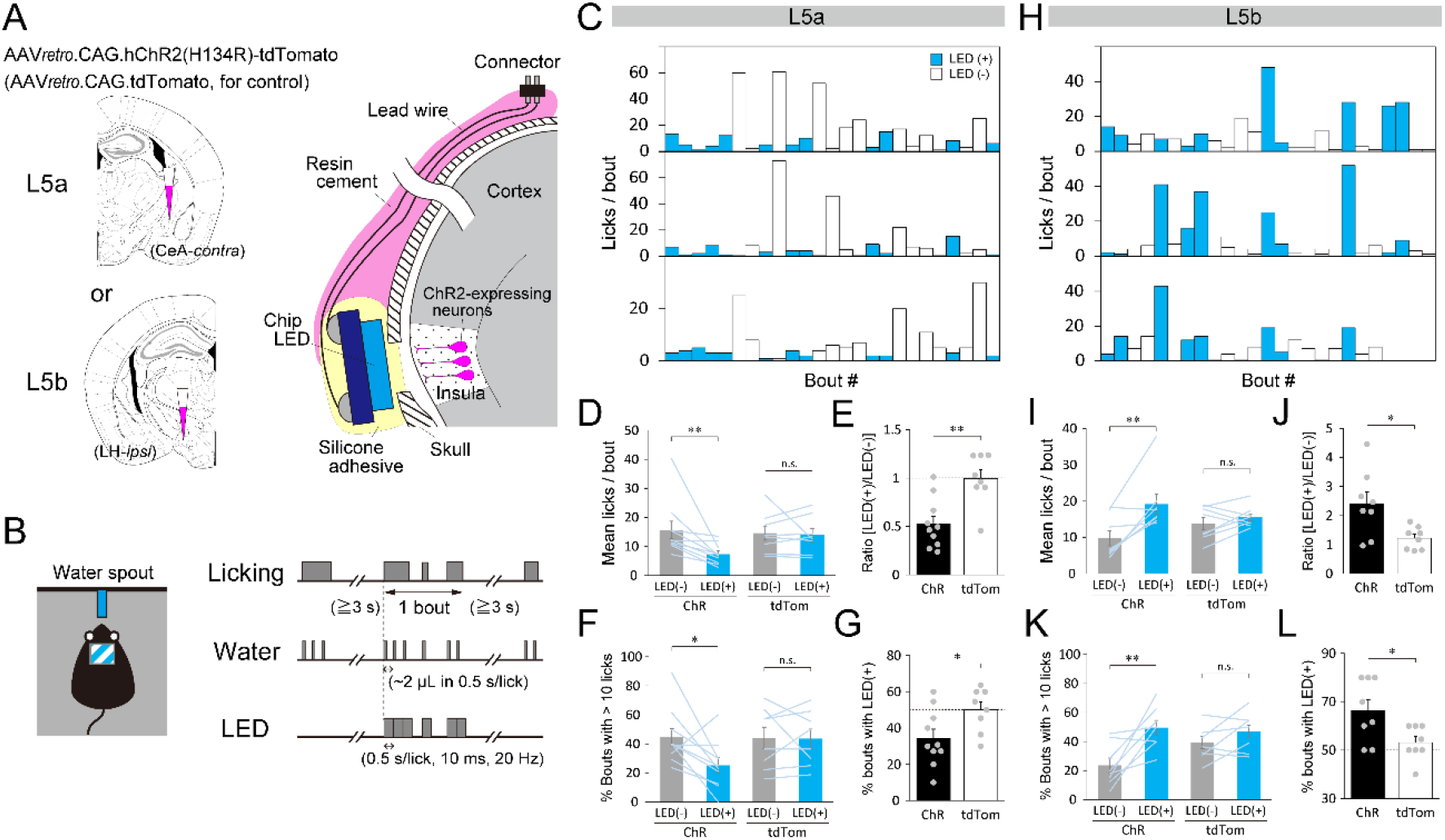
Opposite effects of the optogenetic activation of L5 sublayer neuronal populations on appetitive licking. (A) Illustrations depicting AAV injections for retrograde ChR2 expression in the L5a or L5b populations (left) and the placement of a chip LED (right). (B) An illustration depicting a freely moving mouse with a single water spout (left) and the protocol for water delivery and illumination (right). Water (~2 μl) is delivered when the mouse licks the spout. The illumination (10 ms, 20 Hz) is presented simultaneously at the beginning of water delivery only at random bouts. (C, H) Three typical examples of licking across a series of bouts with (blue) or without (white) illumination for mice expressing ChR2 in the L5a (C) and L5b (H) populations. Only a part of the session (25 bouts) is shown. (D, I) Comparisons of mean licks per bout between the bouts with (blue) and without (gray) LED illumination for ChR2-expressing (ChR) and control (tdTom) mice (L5a: 15.7 ± 3.0 for [LED-] vs. 7.4 ± 1.1 for [LED+], p = 0.0039 for ChR; 14.6 ± 2.5 for [LED-] vs. 13.9 ± 2.2 for [LED+], p = 0.84 for tdTom, D; L5b: 9.7 ± 2.0 for [LED-] vs. 19.2 ± 2.8 for [LED+], p = 0.0063 for ChR; 13.8 ± 1.6 for [LED-] vs. 15.5 ± 1.0 for [LED+], p = 0.30 for tdTom, I). (E, J) Ratio of the mean licks per bout with illumination relative to those without illumination (L5a: 0.53 ± 0.08 for ChR vs. 0.99 ± 0.09 for tdTom, p = 0.0015, E; L5b: 2.41 ± 0.40 for ChR vs. 1.22 ± 0.14 for tdTom, p = 0.022, J). (F, K) Proportion of long drinking bouts with > 10 licks per bout in the session (L5a: 44.6 ± 5.9% for [LED-] vs. 25.2 ± 5.2% for [LED+], p = 0.028 for ChR; 44.0 ± 7.4% for [LED-] vs. 43.5 ± 6.6% for [LED+], p = 0.93 for tdTom, F; L5b: 23.6 ± 5.0% for [LED-] vs. 49.5 ± 5.0% for [LED+], p = 0.0095 for ChR; 39.3 ± 4.0% for [LED-] vs. 46.9 ± 4.5% for [LED+], p = 0.12 for tdTom, K). (G, L) Percentage of bouts with illumination in the top 10 longest bouts in the session (L5a: 34.5 ± 4.9% for ChR vs. 50.0 ± 4.3% for tdTom, p = 0.034, G; L5b: 66.4 ± 4.6% for ChR vs. 53.1 ± 2.5% for tdTom, p = 0.022, L). Lines in D, F, I, and K and dots in E, G, J, and L represent individual mice (L5a: n = 10 for ChR, n = 8 for tdTom; L5b: n = 8 for ChR, n = 8 for tdTom). Dotted lines in E, G, J, and L indicate the chance level. * p < 0.05, ** p < 0.01.

We first tested whether and how licking of a water spout was influenced by the activation of each neuronal population in freely moving mice. To execute this, we presented the LED light during licking in random drinking bouts (a bout is a licking cluster sandwiched by ≥ 3 s non-licking periods) (Fig. 4B). In mice expressing ChR2 in the L5a population, we found that the number of licks was markedly smaller in many bouts with LED illumination than in the bouts without illumination (Fig. 4C). The mean number of licks per bout was significantly smaller in the bouts with illumination compared with those without illumination (Fig. 4D). Because the licking behavior was still observed during illumination (Fig. 4D, 7.4 ± 1.1 licks per bout with illumination), it is unlikely that the illumination directly interrupted licking and swallowing movements. When tested in control mice whose L5a population expressed only tdTomato (Fig. 4A), the mean number of licks per bout showed no change in the bouts with LED illumination compared with those without illumination (Fig. 4D), indicating that the illumination per se has no effect on drinking behavior. The mean number of licks per bout was reduced approximately by half on average by the illumination in the mice with ChR2 expression (Fig. 4E). The suppression of licking by opto-activation of the L5a population was also found in the maximum number of licks per bout (Fig. S4E). Further, the proportion of long drinking bouts (with more than 10 licks per bout) in the session was significantly lower for the bouts with illumination compared with those without illumination, while no difference was found in the control mice (Fig. 4F). In addition, the percentage of bouts with illumination in the top 10 longest bouts (with the highest number of licks) was also significantly lower in the ChR2-expressing mice than in control mice (Fig. 4G). In contrast to the mice whose L5a population expressed ChR2, mice whose L5b population expressed ChR2 showed markedly higher number of licks per bout in the bouts with illumination compared with those without illumination (Fig. 4H, I). The mean number of licks per bout was increased more than twice on average by the illumination in the ChR2-expressing mice (Fig. 4J). The facilitation of licking by opto-activation of the L5b population was also manifested in the maximum number of licks per bout (Fig. S4F), the proportion of long drinking bouts (Fig. 4K), and the percentage of bouts with illumination in the top 10 longest bouts (Fig. 4L).

The behavioral changes induced by opto-activation of the L5 neuron subpopulations are either caused by emotional responses, that is, an avoidance against and a preference for the spout with illumination that led to the suppression and facilitation of licking, respectively, or are attributed only to the modulation of the motivational aspect of licking without emotional change. To address this question, we performed a two-spout choice test in which mice can freely choose either the spout paired with illumination or the unpaired one, both spouts of which deliver equal amount of water per lick (Fig. 5A). We reasoned that if opto-activation during licking induces aversive emotional valence, the mice would show preference towards the spout unpaired with illumination and that if opto-activation induces appetitive emotional valence, the mice would prefer the spout paired with illumination. This argument was supported by our two-spout choice test results using aversive or appetitive tastants in intact B6 mice. The mice preferred the water spout over the quinine spout in the water-quinine choice test and the sucrose spout over the water spout in the water-sucrose choice test while showing no preference between the two spouts delivering water (Fig. S5A). The mice also exhibited more frequent spout switching when they licked the quinine spout in the water-quinine choice test and less frequent spout switching when they licked the sucrose spout in the water-sucrose choice test (Fig. S5B). Testing the mice, whose L5a population expressed ChR2, with both spouts delivering water revealed that opto-activation reduced the mean and maximum number of licks per bout and the proportion of long bouts (Fig. 5C, Fig. S5C-E), in line with the single-spout test results (Fig. 4D-G, Fig. S4E), but affected neither the choice of spout nor spout switching (Fig. 5B). These results suggest that opto-activation of the L5a population has no effect on negative emotional valence but suppresses the motivation of licking.

**Figure 5.**
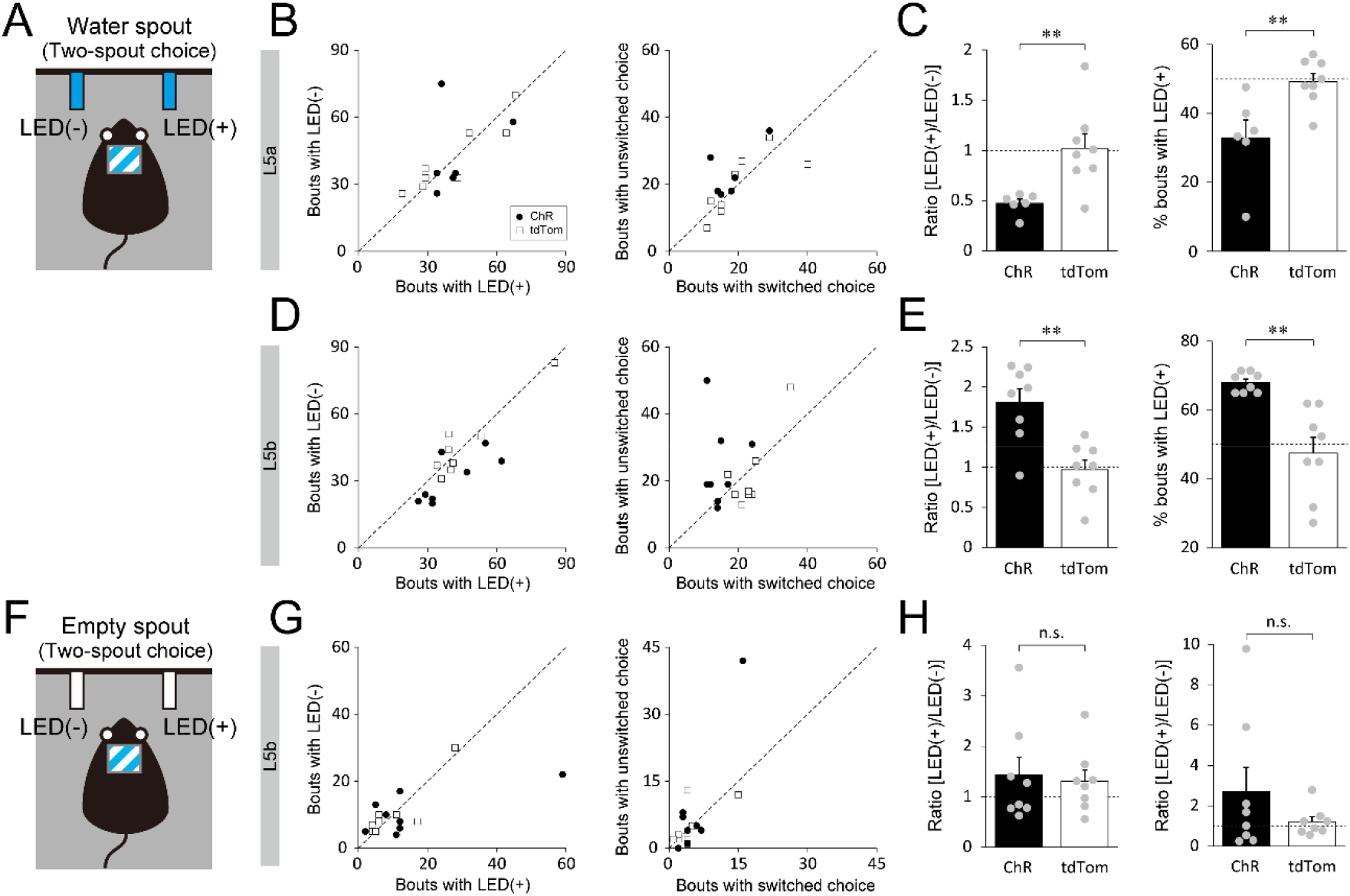
Lack of emotional valence induced by optogenetic activation of L5 subpopulations. (A) An illustration depicting a freely moving mouse in a two-spout choice test with one water spout paired with LED illumination and the other unpaired with LED illumination. Note that the two spouts deliver the same amount of water per lick. (B, D) Left: the number of bouts in which the mouse chose the spout with vs. without illumination for mice expressing ChR2 (black circles) or tdTomato (white squares) in the L5a population (ChR: 42.3 ± 5.1 for [LED+] vs. 43.7 ± 7.7 for [LED-], p = 0.56; tdTom: 41.0 ± 6.3 for [LED+] vs. 41.8 ± 5.4 for [LED-], p = 0.78, B) and L5b population (ChR: 39.9 ± 4.7 for [LED+] vs. 31.3 ± 3.8 for [LED-], p = 0.024; tdTom: 45.9 ± 5.9 for [LED+] vs. 46.1 ± 5.8 for [LED-], p = 0.91, D). Right: the number of bouts in which the mouse switched vs. unswitched choice next to the bouts with illumination (ChR: 17.8 ± 2.5 for switched choice vs. 23.2 ± 3.1 for unswitched choice, p = 0.071; tdTom: 20.3 ± 3.5 for switched choice vs. 19.8 ± 3.2 for unswitched choice, p = 0.84 for the L5a population, B; ChR: 14.8 ± 1.5 for switched choice vs. 24.5 ± 4.4 for unswitched choice, p = 0.047; tdTom: 23.4 ± 1.9 for switched choice vs. 21.8 ± 4.0 for unswitched choice, p = 0.56 for the L5b population, D). (C, E) Left: the ratio of the mean licks per bout with illumination relative to those without illumination for mice expressing ChR2 (black) or tdTomato (white) in the L5a population (0.47 ± 0.04 for ChR vs. 1.02 ± 0.14 for tdTom, p = 0.0062, C) and L5b population (1.81 ± 0.17 for ChR vs. 0.97 ± 0.12 for tdTom, p = 0.0012, E). Right: the percentage of bouts with illumination in the top 10 longest bouts in the sessions (L5a: 33.0 ± 5.2% for ChR vs. 49.3 ± 2.4% for tdTom, p = 0.0085, C; L5b: 68.0 ± 1.0% for ChR vs. 47.5 ± 4.6% for tdTom, p = 0.0026, E). Gray dots represent individual mice (n = 6 for ChR, n = 8 for tdTom in B and C; n = 8 for ChR, n = 8 for tdTom in D and E). (F) An illustration depicting a freely moving mouse in a two-spout choice test with one spout paired with and the other unpaired with LED illumination. Note that both spouts were devoid of water. (G) Left: the number of bouts in which the mouse chose the spout with vs. without illumination for mice whose L5b population expressed ChR2 (black circles) or tdTomato (white squares) (ChR: 15.1 ± 6.4 for [LED+] vs. 10.6 ± 2.2 for [LED-], p = 0.64; tdTom: 10.1 ± 3.0 for [LED+] vs. 10.4 ± 2.9 for [LED-], p = 0.42). Right: the number of bouts in which the mouse switched vs. unswitched choice next to the bouts with illumination (ChR: 5.6 ± 1.6 for switched choice vs. 8.9 ± 4.8 for unswitched choice, p = 0.53; tdTom: 4.6 ± 1.6 for switched choice vs. 5.0 ± 1.7 for unswitched choice, p = 0.88). (H) Left: the ratio of the mean licks per bout with illumination relative to those without illumination for mice whose L5b population expressed ChR2 or tdTomato (1.43 ± 0.35 for ChR vs. 1.31 ± 0.22 for tdTom, p = 0.88). Right: the ratio of the total number of licks in the session (2.70 ± 1.20 for ChR vs. 1.21 ± 0.25 for tdTom, p = 0.88). Gray dots represent individual mice (n = 8 for ChR, n = 8 for tdTom). Dotted lines in B-E, G, and H indicate the chance level.

Activation of the L5b population revealed that a portion of mice showed a higher frequency of choice of the spout with illumination compared with of the spout without illumination and less frequent spout switching when they chose the spout with illumination. On average, the number of choices of the illumination-paired spout and that of unswitched choices next to the bouts with illumination were slightly but significantly higher than that of the illumination-unpaired spout and that of switched choice, respectively (Fig. 5D). Additionally, a marked increase was noted in the mean and maximum number of licks per bout with illumination and the proportion of long drinking bouts with illumination in ChR2-expressing mice (Fig. 5E, Fig. S5F-H). To test whether opto-activation of the L5b population is sufficient to induce a positive valence and elicit the preference for the spout with illumination, using the same group of mice (non-deprived of water before the test), we further performed another two-spout choice test in which one of the two spouts was paired with illumination, but neither delivered water (Fig. 5F). The results showed that the mice did not frequently lick the spout even paired with illumination. There was no significant difference in the number of choices between the spouts with and without illumination and between switched and unswitched choices, although one of the eight ChR2-expressing mice tested exhibited exceptionally a remarkable preference for the spout with illumination (Fig. 5G). Similarly, the mean number of licks per bout and the total number of licks in the session were not significantly increased by illumination in ChR2-expressing mice compared with control mice (Fig. 5H). Therefore, it is unlikely that the activation of the L5b population per se induces emotional valence, although it may enhance the incentive to drink water.

## Discussion

Functional differences among anatomically distinct L5 neuronal subpopulations have been reported in sensory (44, 45), frontal/motor (46, 47), prefrontal (48, 49), and anterior cingulate (50) cortices of rodents. The anatomical and functional heterogeneity in L5 neurons of the insula, however, remains poorly understood, despite the wide variety of subcortical efferents of the insula and their difference between A-P subregions (2, 19, 21, 23). In this study, by taking advantage of the hemispheric difference in the subcortical projections, we succeeded in dissecting L5 neurons of the DgI into two subpopulations with distinct sublaminar distributions, molecular expressions, axonal targets in the extended amygdala and hypothalamus, and modulatory roles in appetitive behavior (Fig. 6).

**Figure 6.**
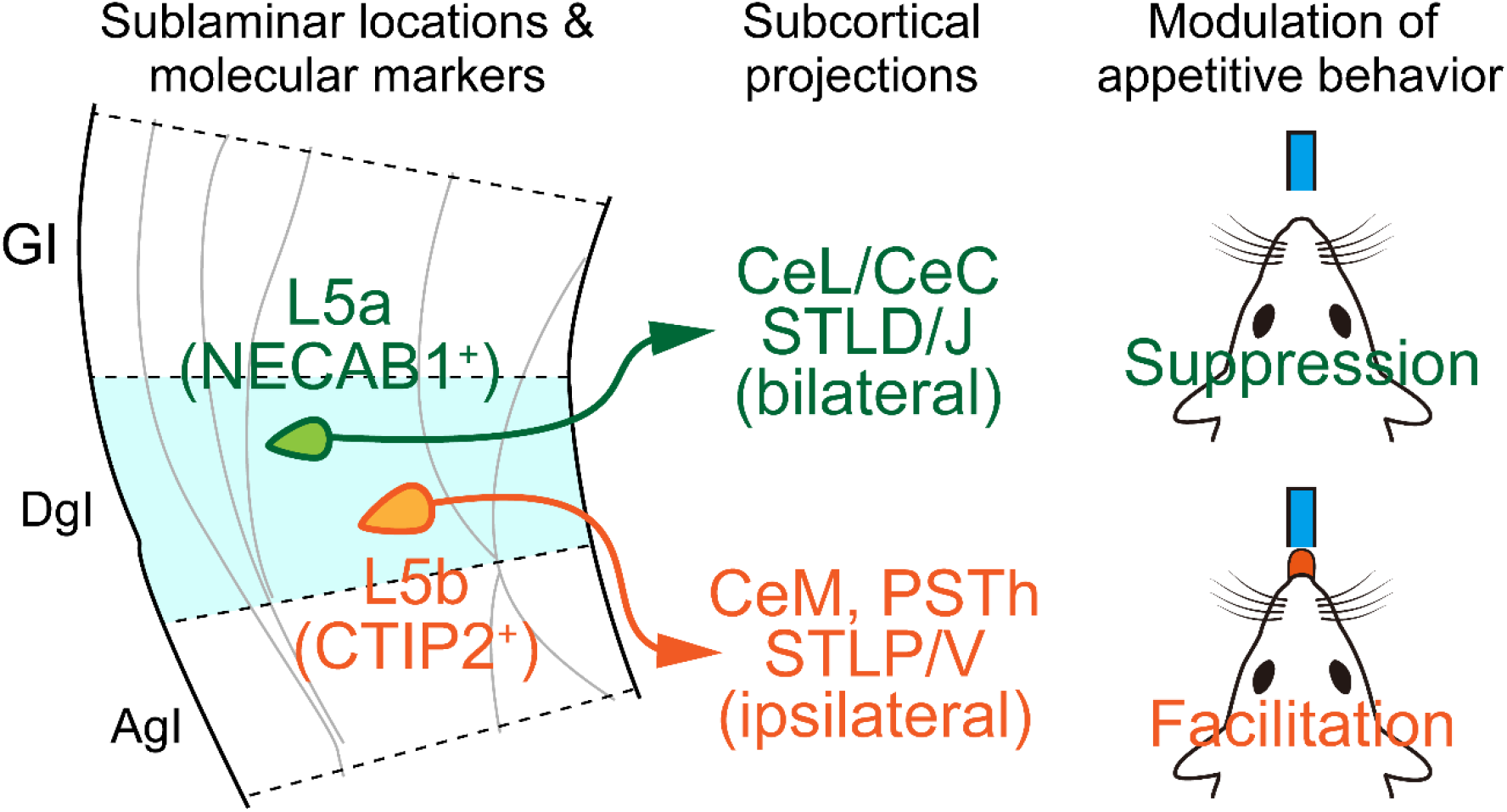
Summary of the study. Molecular, anatomical, and functional characteristics of sublaminar neuronal populations in L5 of the DgI.

### L5a population as a suppressor of appetitive behavior

Previous optogenetic studies have suggested that the insula-CeA circuit serves to suppress feeding and drinking behaviors (2, 3, 8, 14). Because ChR2 was expressed in the insula ipsilateral to the illuminated side in all of the studies, it is plausible that the illumination above ChR2-expressing axon terminals in the CeA simultaneously activated the axons originating from multiple layers of the insula, including the L5a subpopulation (projecting to the CeL/CeC) and L5b subpopulation (projecting to the CeM), which contribute opposingly to appetitive drinking, based on our results (Fig. 4). In this sense, the L5a to CeL/CeC pathway is likely a critical component of insula-CeA circuits involved in the suppression of appetitive behavior. Possibly, the L5a to CeL/CeC pathway was activated more effectively than the L5b to CeM pathway by optogenetic techniques used in the previous studies. It is also possible that the activity of the L5a-CeL/CeC circuit has more impact on appetitive behavior by default when both L5a and L5b pathways are activated simultaneously. Nevertheless, it is worth noting that the net effect of insula L5-CeA circuit activities on appetitive behaviors depends on how strong each of the sublayer axons is synaptically connected with functionally diverse subpopulations of CeA neurons and on which of the connections is potentiated by experiences (14).

Our evidence is against the interpretation of licking suppression as an aversive response because opto-activation of the L5a population induced no avoidance against the water spout paired with illumination in our two-spout choice test. This result contradicts that from previous studies showing that optogenetic activation of the insula (1, 3, 4) or insula-CeA circuits (2, 8) induced an avoidance behavior in a place preference test, although no anxiogenic effect was observed in an open-field test with opto-activation of insula-CeA circuits (14). It is possible that the L5a population merely serves to attenuate the incentive salience in appetitive behavior, and other neuronal populations and their circuits are involved in the generation of negative emotional valence. One may wonder whether the licking suppression by opto-activation of the L5a population was due to the interruption of licking and swallowing movements rather than the attenuation of the motivation. Our observation of licking behavior during the illumination, despite the number was smaller than that in the bouts without illumination, argues against this possibility (Fig. 4 C-G). Moreover, if the mice had been interrupted from drinking by illumination in the two-spout choice test, they should have preferred the spout without illumination; but the result showed no preference for the spout (Fig. 5B). It is thus most likely that the opto-activation suppressed the motivational aspect of drinking behavior, possibly inducing transient satiety in the bouts with illumination.

### Possible neural pathways of L5b involved in the facilitation of appetitive behavior

In contrast to the suppressive effect by the L5a population, we found the facilitatory effect of the L5b opto-activation on drinking behavior (Fig. 4H-L) but not on licking behavior per se (Fig. 5H), suggesting that the L5b population may contribute to the enhancement of incentive salience by recruiting ‘wanting’ neural circuits (51). This notion is in line with the functional properties of the CeM, the amygdala target region of the L5b population, where essentially all neuron subtypes promote appetitive behaviors (35). However, the role of the PSTh, another target of the L5b population, is currently controversial; it responds to palatable food (29) but is activated in neophobic (42) and anorexigenic (40, 42) situations and is also activated by a conditioned aversive taste although not by bitter taste (39). The latter results agree with a study in which optogenetic activation of a PSTh neuronal population projecting to the PVT resulted in feeding suppression (41). The controversy in the role of PSTh could be attributable to its neuronal diversity; the PSTh contains cell populations expressing preprotachykinin-1 (29, 42, 52), calbindin (29, 53), corticotropin-releasing hormone (40), or vesicular glutamate transporter 2 (41). Therefore, it is crucial to identify the neuronal population of the PSTh that is specifically innervated by the L5b population of the insula. Besides the CeM and PSTh, the L5b population of the insula characterized in our study has multiple subcortical targets, such as the pons and medulla. Insula L5b population is, therefore, reminiscent of pyramidal tract-type neurons in the neocortex (54–56). This idea is supported by the expression of the Ctip2 gene in both neuron types (32, 57; our current results). How the divergent projections of the L5b population contribute to the facilitation of appetitive behavior needs to be unraveled in future studies.

Like the activation of the L5a population, opto-activation of the L5b population in the middle-posterior DgI had no modulatory effect on the emotional aspect of drinking behavior (Fig. 5F, G). However, because the posterior insula has been reported to have a hedonic ‘liking’ hot spot (51, 58), other layers or subregions (e.g., GI and AgI) of the posterior insula might be linked to neural circuits involved in positive emotions.

### A perspective of the insula’s role in taste quality-independent modulation of feeding

Our result of the anterior-low/posterior-high distribution of the L5a population projecting to the CeL/CeC (Fig. 2H) is in line with the previous studies demonstrating an A-P region difference in the insula-mediated modulation of feeding (1) and neural circuits of the anterior insula-BLA and posterior insula-CeA involved in facilitation and suppression, respectively (2). Our finding of the distribution of the L5b population projecting to the PSTh/CeM with an anterior-high/posterior-low gradient (Fig. 2H) further suggests its participation in the facilitatory role of the anterior insula (1) in concert with the neuronal populations connecting to the BLA (2). In contrast, despite the difference in distributions along the A-P axis, both L5 subpopulations in the DgI are found along the *entire* axis. Considering that taste quality is represented topographically across the gustatory insula, although some of the neurons can respond to multiple taste qualities (15, 59; cf., 60, 61), our results provide a potential neural substrate for taste quality-independent modulation of feeding rather than taste quality-specific hard-wired modulatory circuits. An issue to be addressed in the future is whether the activation of the L5b subpopulation, as a neural substrate for taste-independent facilitation of feeding, can overcome aversive tastes, although it has been reported that optogenetic activation of the posterior insula-CeA circuit suppresses the intake of an appetitive taste (2, 3). In addition to by taste qualities, motivation for feeding can be affected by a variety of contexts, including homeostatic, emotional, and external circumstances, all of which can be associated with any type of taste by learning. Thus, clarifying the properties of sensory and contextual inputs to each L5 sublayer of the insula is also a critical issue for future studies. Additionally, it is intriguing to address whether the insula L5 sublayers contribute to the modulation of not solely feeding behavior but rather motivational behaviors in general.

## Acknowledgments

We thank Karl Deisseroth for providing pAAV-Ef1a-mCherry-IRES-Cre, Karel Svoboda for providing AAV-CAG-hChR2-H134R-tdTomato, and Edward Boyden for providing pAAV-CAG-tdTomato. We also thank Drs. Junichi Nabekura and Nobuhiko Yamamoto for helpful comments on the early version of this manuscript.

## Funding

This work was supported by the following grants:

JSPS KAKENHI Grant Number 20K06928 (M.T.)

JSPS KAKENHI Grant Number 19K06908 (W.J.S.)

JSPS Grant-in-Aid for Scientific Research on Innovative Areas “Adaptive Circuit Shift” (15H01442) and “Multi-scale” (19H05222) (W.J.S.)

Takeda Science Foundation (M.T.)

This work was also supported by Messrs. Toshihiro Morikawa, Daisuke H. Tanaka, and 34 crowdfunding backers on the *“academist”* platform (https://academist-cf.com/projects/91). (M.T.)

## Author Contributions

Conceptualization, M.T.; Methodology, M.T., S.K., K.K., and W.J.S.; Investigation, M.T.; Supervision, W.J.S.; Writing—Original Draft: M.T.; Writing—Review & Editing: M.T., K.K., and W.J.S.; Funding Acquisition: M.T. and W.J.S.

## Competing Interest Statement

The authors declare no competing interests.

## STAR Methods

### Animals

Male C57BL/6J (B6) mice (Japan SLC, Shizuoka, Japan) were used for all experiments. At the time of the injection experiments (see below), the mice were 6-10 weeks old for anatomical studies and were 5-7 weeks old for optogenetic studies. All experiments were approved by the Committee for Animal Experiments of Kumamoto University and performed in accordance with the Guidelines for Use of Animals in Experiments of Kumamoto University.

### Surgery

Mice were anesthetized with a mixture of ketamine (Ketamine injection, Fujita Pharmaceutical, Tokyo) and xylazine (Selactar, Bayer Yakuhin, Tokyo) (ketamine 80 mg/kg, xylazine 8 mg/kg) during surgery. For injections of CTB or viruses into subcortical regions (i.e., the CeA and PSTh), mice were head-fixed on a stereotaxic frame (SR-5M-HT, Narishige, Tokyo). After partial hair removal and incision of the skin along the midline of the head to expose the skull, two small craniotomies (~1 mm square for each) were made over the locations of the injection (one for the right CeA, the other for the left PSTh). After the injections, the parietal skin was sutured and glued with a tissue adhesive (Vetbond, 3M, St. Paul, MN). For virus injections into the insula, a mouse was head-fixed on a rotatable head holder (SG-4N, Narishige, Tokyo). After rotating the head 70-80 degrees, the skin of the temporal head (left side) was incised and muscle tissues were peeled to expose the skull. A small craniotomy was performed over the middle-posterior insula at the midpoint between the middle cerebral artery and the vertical squamosal suture immediately dorsal to the ventral eminence of the squamosal bone (62). For LED implantations, the craniotomy was performed in the same way as for virus injections into the insula, except for the area of skull removal being a ~ 0.8 × 1.2 mm rectangle. Subsequently, a blue tip LED (470 nm, OSBL1608C1A, OptoSupply, Hong Kong) connected with lead wires, the soldered portions of which were insulated with modified silicone resins, was placed over the cranial window so that the LED surface was faced to the exposed cortical surface. The LED was then covered with a silicone adhesive (Kwik-Sil, World Precision Instruments, Sarasota, FL) together with the peeled tissues and skin and further covered with the Kwik-Sil and Vetbond. Using a resin cement (Super-Bond, Sun Medical, Shiga, Japan), the lead wires were fixed on the skull, and a connector was placed on the parietal portion of the skull (see Fig. 4A) with a screw embedded into the frontal part of the skull. Immediately after surgery, the mice were injected subcutaneously with buprenorphine (0.1 mg/kg of the body weight, Lepetan, Otsuka Pharmaceutical, Tokyo) and enrofloxacin (5 mg/kg of the body weight, Baytril, Bayer Yakuhin, Tokyo) for postoperative analgesia and blocking of bacterial infections, respectively. All mice were individually housed and maintained in a breeding room on a 12 h light-dark cycle with food and water available ad libitum. The mice were housed there for 3 days after CTB injections or 2 weeks after virus injections for axon tracing until sacrifice, and for 5-7 weeks after virus injections for optogenetics and at least for another 5 days after LED implantations until behavioral experiments.

### Tracer and virus injections

All injections of tracers and viruses were performed by air pressure (0.05 μl per 1-3 min) with a custom-made syringe pump connected to a glass capillary needle (Drummond precision calibrated micropipettes #2-000-001, ~40 μm tip diameter) filled with injection solutions. For labeling axonal projections of the insula, an adeno-associated virus (AAV) vector (AAV2-CAG-GFP or AAV2-CAG-FLEX-GFP, University of North Carolina Vector Core, 0.2 μl) was injected 0.4 mm deep from the surface of the middle-posterior insula (0.05 ± 0.3 mm posterior to Bregma, mean ± standard deviation (SD), n = 9 mice: 3 mice for AAV2-CAG-GFP, 6 mice for AAV2-CAG-FLEX-GFP). For retrograde tracing, the CTB conjugated with Alexa Fluor 488 (CTB-green) or 555 (CTB-red) (Invitrogen, approximately 0.05 μl of 5 mg/ml dissolved in 0.1 M phosphate buffer) (63) was injected into the CeA (1.5 mm posterior, 3.0 mm lateral, 4.5 mm ventral to Bregma, but 0.8 mm posterior, 2.5 mm lateral, 4.9 mm ventral to Bregma for localized injections into the CeM) or the PSTh (2.0 mm posterior, 1.1 mm lateral, 5.0 mm ventral to Bregma). For the mice aged less than 7-week-old, the coordinates of the injection sites were 0.1 mm smaller than the Bregma levels described above. For sublayer-specific protein expressions (i.e., the Cre-inducible GFP, ChR2-tdTomato, and tdTomato), retrograde AAVs (34) described below were injected into either the right CeA or left PSTh. The pAAV-Ef1a-mCherry-IRES-Cre was a gift from Karl Deisseroth (Addgene viral prep # 55632-AAVrg; http://n2t.net/addgene:55632; RRID:Addgene_55632) (64). The AAV-CAG-hChR2-H134R-tdTomato was a gift from Karel Svoboda (Addgene viral prep # 28017-AAVrg; http://n2t.net/addgene:28017; RRID:Addgene_28017) (65). The pAAV-CAG-tdTomato (codon diversified) was a gift from Edward Boyden (Addgene viral prep # 59462-AAVrg; http://n2t.net/addgene:59462; RRID:Addgene_59462).

### Immunohistochemistry

To identify the CeL/CeC, we performed immunohistochemistry against PKC-δ (30) or the regulator of G protein signaling 14 (RGS14) (66–68) as the restricted pattern of RGS14 immunoreactivity to the CeL/CeC was observed in the present study (Fig. 3D, H). For cortical layer markers, immunohistochemistry against the forkhead box protein P2 (FOXP2) for L6 (69, 70), NECAB1 for L5a (31), and CTIP2 for L5b (32, 33) was performed. Sections were incubated in PBS containing 0.5% normal donkey serum and 0.03% Triton X-100 for 1 h at room temperature (RT, 23 ± 7°C) and then stained with primary antibodies overnight at 4°C (rabbit anti-c-Fos, 1:5000, Sigma-Aldrich; rat anti-CTIP2, 1:1000, Abcam; rabbit anti-FOXP2, 1:1000, Abcam; mouse anti-FOXP2, 1:2000, Atlas Antibodies; rat anti-GFP, 1:2000, Nacalai; rabbit anti-NECAB1, 1:2000, Atlas Antibodies; mouse anti-PKC-δ, 1;500; mouse anti-RGS14, 1:500, NeuroMab; two overnight incubations for c-Fos and a triple-staining of NECAB1, CTIP2, and FOXP2). After washing three times in PBS for 10 min, the sections were incubated with secondary antibodies for 2 h at RT (donkey anti-mouse IgG Alexa-555 conjugate for PKC-δ and FOXP2; donkey anti-mouse IgG Alexa-647 conjugate for RGS14; donkey anti-rabbit IgG Alexa-488 conjugate for c-Fos; donkey anti-rabbit IgG Alexa-647 conjugate for c-Fos and FOXP2; donkey anti-rabbit IgG Alexa Plus-647 conjugate for NECAB1; donkey anti-rat IgG Alexa-488 conjugate for GFP and CTIP2; donkey anti-rat IgG Alexa Plus-647 conjugate for CTIP2; 1:1000 dilution for all the secondary antibodies, Invitrogen). After washing three times in PBS for 10 min, the sections were mounted on glass slides and counterstained with DAPI. Coverslips were finally mounted with 10% Mowiol 4-88 (Polysciences, Warrington, PA) dissolved in 0.1 M Tris-HCl (pH 8.0) and 25% (w/v) glycerol. The fluorescent images were taken through a confocal laser scanning microscope (FV3000, Olympus, see below) or a fluorescence microscope (BZ-X800, Keyence, Osaka, Japan).

### Optogenetics

The LED (see Surgery) was controlled wirelessly via a receiver unit (TeleR-1-P, TeleOpto, Bio Research Center, Nagoya, Japan) and an infrared emitter (TeleEmitter, Bio Research Center, Nagoya, Japan) connected to a remote controller (TeleRemocon, Bio Research Center, Nagoya, Japan). The controller was operated through a microcontroller (Arduino UNO, Arduino, Somerville, MA) with a custom-made program (on the Arduino IDE). The illumination was a train of 10 pulses (10-ms pulse duration and 50-ms interpulse interval) and was delivered in response to a lick (see Fig. 4B). The maximum LED powers (measured before their implantation) were 10 to 20 mW per LED (0.96 mm2). For the confirmation of effective optogenetic activations, c-Fos expression in the insula was examined for all mice expressing ChR2 used in the behavioral experiments. To do this, within two weeks of the end of a series of behavioral experiments, the mice were deeply anesthetized and perfused with 4% PFA after repetitive illuminations of the LED for 90 min in a plastic cage (2 s trains of 10 ms pulses at 20 Hz every 4 s), and the brain sections were immunostained with the anti-c-Fos antibody (see Immunohistochemistry).

### Behavioral experiments

To connect a wireless receiver unit immediately before starting a session (including habituation sessions), mice were weakly anesthetized with 4% isoflurane for 2-3 min and then maintained with 2% isoflurane for a minute while the receiver unit was connected to the connector on the head. The mice were then recovered in a home cage for > 5 min. Habituation sessions were performed for 3 h on two consecutive days. During the session, the mice (not deprived of water before the sessions) were put in a plastic cage (W 16.5 cm × D 23 cm × H 12 cm, maximum inside dimensions) equipped with a single water spout (Feeding needle KN-348-20G, Natsume Seisakusho, Tokyo) at the height of 4.5 cm from the bottom of the cage, and the cage was placed in a sound-attenuating chamber. Through the spout, a drop of water (approximately 2 μl) was delivered every time the mice licked the spout ad libitum. No water was available for 0.5 s after each water drop. The mice were returned to the home cage after the habituation sessions and deprived of water for ~18 h before starting the test session. A single-spout test session was first carried out for 90 min in which the LED illumination pulse train was delivered in response to a lick in approximately half of the drinking bouts in the session. An interruption of continuous licking for 3 s or longer was considered as the end of a drinking bout. The bout to be LED illuminated was determined in a pseudorandom order. Water was delivered in the same way as in the habituation session regardless of the presence or absence of LED illuminations. Within a week after the single-spout test, two-spout choice tests were performed in which the water-deprived (18 h) mice were put in another plastic cage (W 20 cm × D 31 cm × H 13 cm, maximum inside dimensions) equipped with two water spouts (~14 cm apart from each other), which delivered equal amount of water (~2 μl) per lick. A 60 min daily session was performed for two consecutive days. The LED illumination was paired with water delivery from the right spout in the first session, and the pairing was switched to the left spout in the second session to minimize potential spatial bias. Mice with their L5b population expressing ChR2 or tdTomato received a third session (60 min) of two-spout choice test within three days after the second two-spout session. In the third session, the mice were not deprived of water before the test, and the two spouts, one of which (the right side) was paired with the LED illumination, delivered no water upon licking.

For taste-biased two-spout choice control tests, one day after the 3-h habituation in a cage equipped with two water spouts, water-deprived (18 h) B6 intact mice were tested with one spout delivering water and the other delivering equal volume of 0.02% quinine hydrochloride in a 60-min session. Two days later, the same mice (18 h water-deprived) were subjected to a second 60-min session in which one spout delivered water and the other delivered equal volume of 2% sucrose. Seven days after the second session, the mice were further subjected to a third session with both spouts delivering water.

### Quantitative analyses

To estimate the A-P level of virus injection sites in the insula, serial coronal sections were obtained from each virus-injected brain (n = 3 mice for AAV2.CAG.GFP injections; n = 6 mice for AAV2.CAG.FLEX.GFP injections), and the Bregma level of the section exhibiting the strongest fluorescence was determined by reference to the mouse brain atlas (71).

To examine the laminar distribution of CTB-labeled cells in the DgI, fluorescent (RGB) images of the coronal sections (the A-P level was between 0.1 mm anterior and 0.2 mm posterior to Bregma) from brains with dual CTB injections (n = 5 mice for CeA-contra/PSTh-ipsi; n = 4 hemispheres from 3 mice for CeA-contra/CeL/CeC-ipsi; n = 4 mice for CeA-contra/CeM-ipsi) were obtained with a confocal laser scanning microscope (10× objective lens, FV3000, Olympus, Tokyo), and z-stack images were created by a maximum intensity projection (ImageJ, NIH) from three confocal slices at 4-μm intervals. Using the ImageJ software, a rectangular region of interest (ROI) (150 μm for the short axis, the long axis was in the direction of cortical depth) was selected in the DgI so that the ROI covered the laminar structure (L1-L6) and the long axis was perpendicular to the border between L5b and L6, indicating cortical depth (layer and subregional boundaries were delineated based on DAPI staining or FOXP2 immunostaining for the L5b-L6 border) (Fig. 1A, Fig. S1D). The green and red components were then extracted separately as grayscale images from the ROI of the original RGB image. Subsequently, a signal intensity profile along the long axis was obtained by averaging the pixel intensity of the grayscale image across the ROI short axis (150 μm, corresponding to 120 pixels). The fluorescence profile in the L2 to L5b range was extracted, and the values in the profile were scaled to between 0 and 1. Finally, the profile was divided into 20 bins (15-21 pixels per bin except for the last bin with 16-28 pixels), and the bin averages are shown in graphs (Fig. 2D, Fig. S2B, C). For quantification of CTB-labeled CeA-contra projector and PSTh-ipsi projector cells along the A-P axis of the insula, fluorescent confocal images (z-stack) were obtained from sections at eight different A-P levels (1.1 ± 0.1 mm, 0.8 ± 0.1 mm, 0.5 ± 0.1 mm, 0.2 ± 0.1 mm anterior to Bregma; 0.1 ± 0.1 mm, 0.4 ± 0.1 mm, 0.7 ± 0.1 mm, 1.0 ± 0.1 mm posterior to Bregma), and the number of CTB-labeled cell bodies in the insula was counted for each section (the GI and L6 of the DgI to AgI were excluded from the analysis). The fraction (%) of labeled cells at each A-P level was calculated by the number of labeled cells at each A-P level divided by the sum of labeled cells for all eight levels. Six and five brains were analyzed for CeA-contra projector cells and PSTh-ipsi projector cells, respectively. For the expression of sublayer marker proteins (NECAB1 and CTIP2) in the CTB-labeled cells, fluorescent confocal images (z-stack images created from ten confocal slices at 2-μm intervals) were obtained from coronal sections from brains with dual CTB injections (CeA-contra/PSTh-ipsi, n = 3 mice) stained for either NECAB1 or CTIP2. For each molecular marker, the proportion of the total number of CTB/marker double-positive cells to CTB-labeled cells within the DgI of three proximal sections between 0.2 mm anterior and 0.5 mm posterior to Bregma (L6 was excluded) was calculated.

For the quantification of c-Fos immunoreactive cells after optogenetic activation of the L5 subpopulation, fluorescent confocal images (single slices) were obtained from both opto-stimulated and non-stimulated side of the insula of a coronal section. To isolate significant c-Fos immunoreactive particles using the ImageJ, a binarized image was created from a grayscale image of the c-Fos immunofluorescence as follows: First, a background image was created from the original grayscale image (“Rolling ball radius” of 50 pixels), and the mean gray value and SD of a 400 μm square ROI (the position of which was arbitrarily selected within the region corresponding to the insula) were measured as the background level. The original grayscale image was then binarized by the threshold of the mean + 20 × SD of the background ROI. The binarized image was further filtered with a median filter (radius = 2 pixels), and particles of 20 μm2 or larger were regarded as significant c-Fos immunopositive particles. Overlapping particles were identified by reference to the corresponding DAPI image and counted separately. The total number of c-Fos immunopositive particles within the DgI L5 of two proximal sections was counted in each hemisphere as the number of c-Fos-positive cells.

For the quantification of licking behavior, the number of licks was counted for each drinking bout across a session, and the mean and maximum licks per bout (and their ratios between bouts with and without LED illumination), the percentage of bouts with > 10 licks, and the percentage of bouts with LED illumination in the top 10 longest bouts were obtained offline. For two-spout choice tests, in addition to the same analyses as for the single-spout test, counts were also compared between bouts with and without illumination, and also between switched and unswitched choices next to the bouts with illumination.

### Statistical Analysis

Statistical analyses were performed using R (version 4.0.4, the R Foundation). For multiple comparisons of the fraction of CTB-labeled cells along the A-P axis, one-way ANOVA followed by Dunnett’s post hoc test was used. The means of the maximum fractions (0.4 ± 0.1 mm posterior to Bregma for the L5a population; 1.1 ± 0.1 mm anterior to Bregma for the L5b population) were used as the control group in Dunnett’s test. For optogenetic and behavioral experiments, the Shapiro-Wilk test was used to test the normality of each data set (the difference between pairs was tested for pairwise comparisons). F-test was further used to test if the variances of two unpaired groups were equal. For comparisons between two unpaired groups, Student’s t-test was used for groups with both normal distributions and homogeneity of variances, Welch’s t-test was used for groups with normal distributions but no significant equality of variances, and Wilcoxon rank sum (Mann–Whitney U) test was used for pairs involving one or two non-normally distributed groups. For comparisons between paired groups, paired t-test was used if the difference values between groups were normally distributed; otherwise, Wilcoxon signed rank test was used. P < 0.05 was considered statistically significant. Data are expressed as mean ± standard error of the mean unless otherwise stated.

Detailed information on the materials used in this study is listed in Table S1.

## Notes

### Competing Interest Statement

The authors have declared no competing interest.

### Summary of Updates

The title and abstract have been slightly modified.

