## Supplementary Materials for "Dissection of insular cortex layer 5 reveals two sublayers with opposing modulatory roles in appetitive behavior"

**This PDF file includes:**

Figures S1 to S5
Table S1

**Other supplementary materials for this manuscript include the following:**

Dataset S1

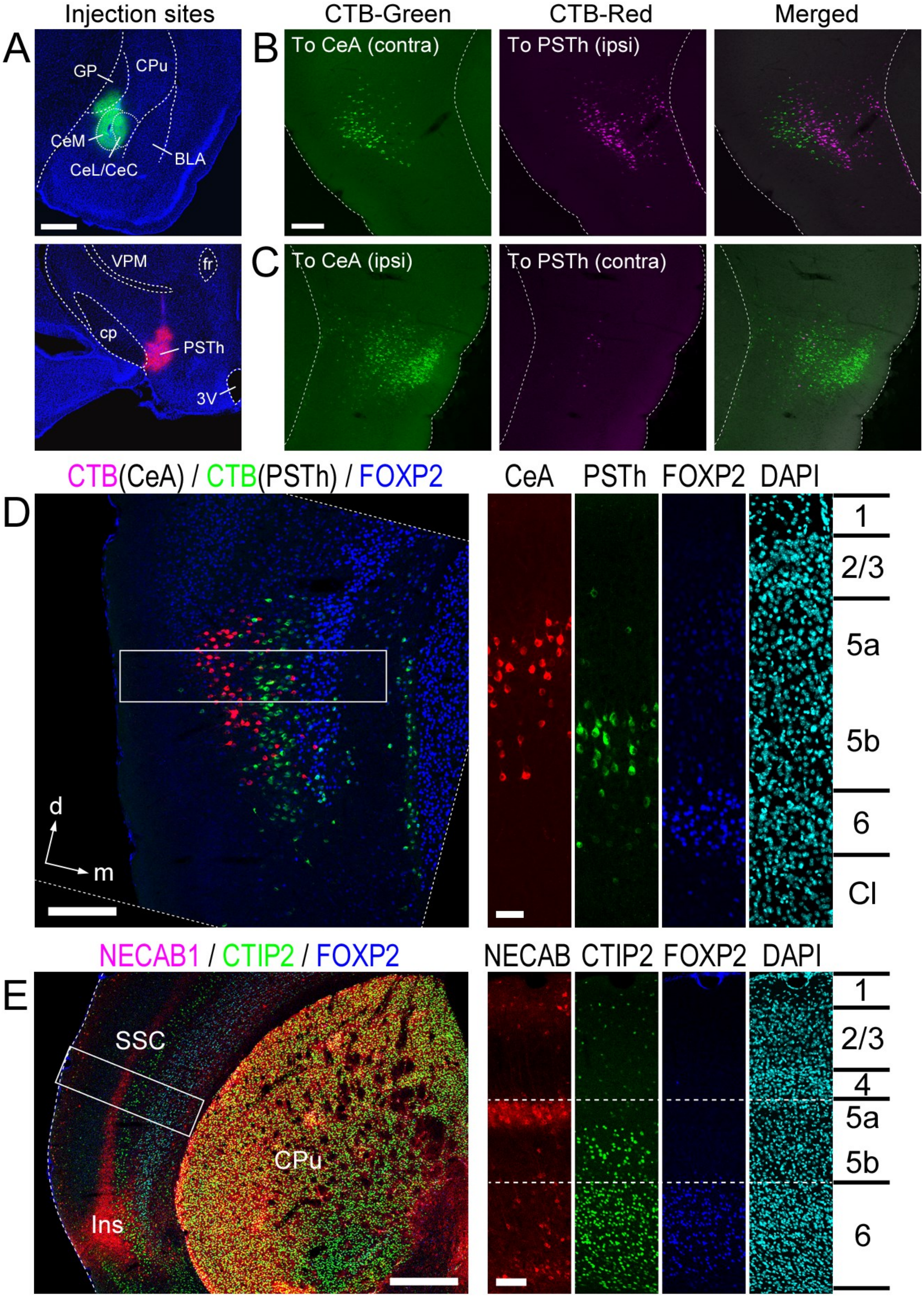

Injection sites
CTB-Green
CTB-Red
Merged
A
GP
CPu
CeM
CeL/CeC
BLA
B
To CeA (contra)
To PSTh (ipsi)
VPM
fr
cp
PSTh
3V
C
To CeA (ipsi)
To PSTh (contra)
CTB(CeA) / CTB(PSTh) / FOXP2
CeA
PSTh
FOXP2
DAPI
D
d
m
1
2/3
5a
5b
6
Cl
NECAB1 / CTIP2 / FOXP2
NECAB
CTIP2
FOXP2
DAPI
E
SSC
CPu
Ins
1
2/3
4
5a
5b
6

**Fig. S1. Distribution of insula cells retrogradely labeled by dual CTB injections into the right CeA and the left PSTh.** (A) Typical example images of the injection sites in the right CeA (top, CTB-green) and the left PSTh (bottom, CTB-red). Sections were counterstained with DAPI (blue). BLA, basolateral amygdala; cp, cerebral peduncle; CPu, caudate-putamen; fr, fasciculus retroflexus; GP, globus pallidus; VPM, ventroposterior medial nucleus of the thalamus, the medial part; 3V, third ventricle. (B) Typical example images of the left insula containing CeA-contra-projecting (green) and PSTh-ipsi-projecting (red) CTB-labeled cells. The left and middle images are the same as shown in Fig. 2C (left and middle), and the right image is a merged one. (C) Typical example images of the right insula from the same brain as in B containing CeA-ipsi-projecting (green) and PSTh-contra-projecting (red) CTB-labeled cells. Note that CeA-ipsi-projecting labeled cells (C, green) were distributed more broadly than CeA-contra-projecting labeled cells (B, green) and that only a few PSTh-contra-projecting labeled cells (C, red) were seen. (D) Sublayer localization of CeA-contra-projecting (red) and PSTh-ipsi-projecting (green) CTB-labeled cells in L5 of the DgI. The left image was rotated clockwise at 14 degrees (dotted lines). The right images are higher magnification views of the box in the left merged image shown in separated fluorescent colors and DAPI staining (cyan). The border between L5 and L6 can be delineated based on the difference of DAPI-stained cell density and on immunostaining for an L6 marker, FOXP2 (blue). Cl, claustrum; d, dorsal; m, medial. (E) Layer-specific expressions of NECAB1 (red), CTIP2 (green), and FOXP2 (blue) in the somatosensory cortex (SSC) and insula (Ins). The right images are higher magnification views of the boxed region (a part of the SSC) in the left merged image shown in separated fluorescent colors and DAPI staining (cyan). Scale bars: 0.5 mm (A and E-left), 0.2 mm (B and D-left), 0.1 mm (E-right), 50 μm (D-right).

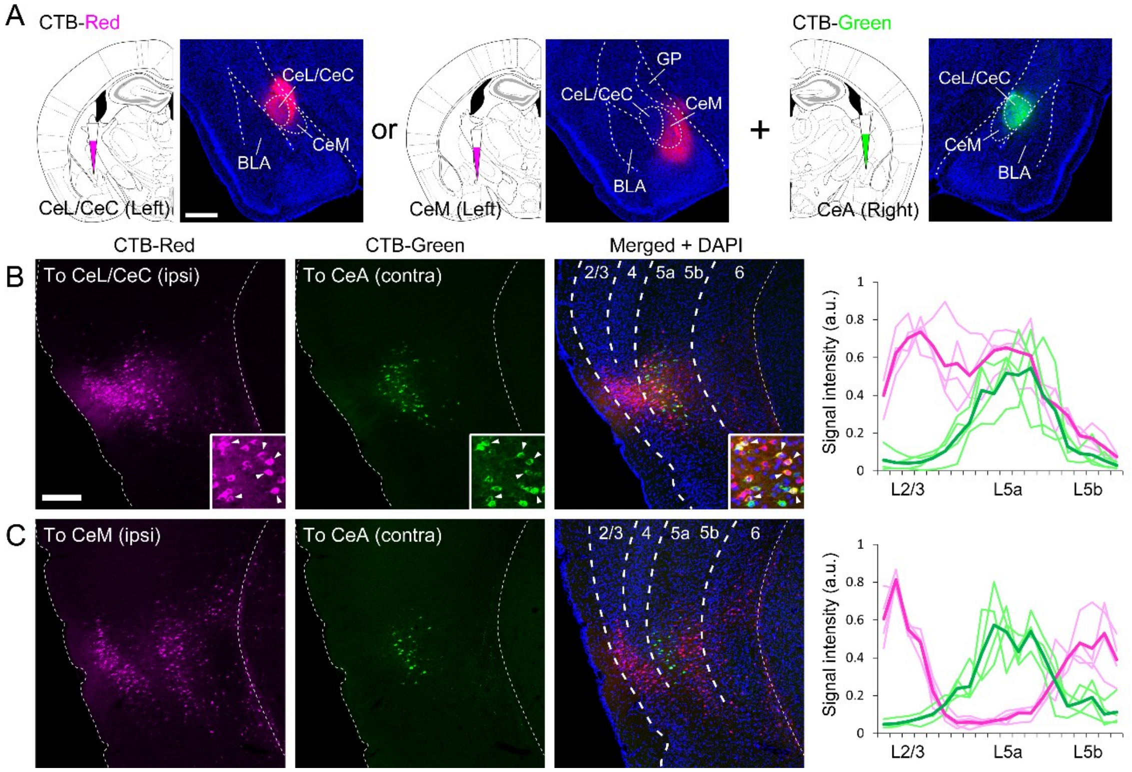


**Fig. S2. Layer-specific DgI projections to the CeA.** (A) Illustrations depicting the injection of CTB-red preferentially into the CeL/CeC or CeM of the left hemisphere, accompanied by the injection of CTB-green into the right CeA. The fluorescent images are typical examples showing the CTB injection sites (counterstained with DAPI) corresponding to the adjacent illustrations. (B) Typical example images of the left insula containing CeL/CeC-ipsi-projecting (red) and CeA-contra-projecting (green) CTB-labeled cells. The right image shows the merged fluorescence (counterstained with DAPI). Double-labeled cells can be seen in L5a (arrowheads in the insets). Right: relative fluorescent intensities (arbitrary unit) of CTB labeling across cortical depth in the DgI (L2/3 to L5b). The signal distributions of CTB-red-labeled cells (projecting to the CeL/CeC-ipsi) and CTB-green-labeled cells (projecting to the CeA-contra) are shown in magenta and green, respectively. Light colors represent individual mice (n = 4 mice with dual CTB injections), and dark colors are the average. Note that peaks are found in L2/3 and L5a for CeL/CeC-ipsi-projecting cells (magenta). (C) Typical example images of the left insula containing CeM-ipsi-projecting (red) and CeA-contra-projecting (green) CTB-labeled cells. The right image shows the merged fluorescence (counterstained with DAPI). Right: relative fluorescent intensities (arbitrary unit) of CTB labeling across cortical depth in the DgI (L2/3 to L5b). The signal distributions of CTB-red-labeled cells (projecting to the CeM-ipsi) and CTB-green-labeled cells (projecting to the CeA-contra) are shown in magenta and green, respectively. Light colors represent individual mice (n = 4 mice with dual CTB injections), and dark colors are the average. Note that peaks are found in L2/3 and L5b but not in L5a for CeM-ipsi-projecting cells (magenta). Scale bars: 0.5 mm (A), 0.2 mm (B).

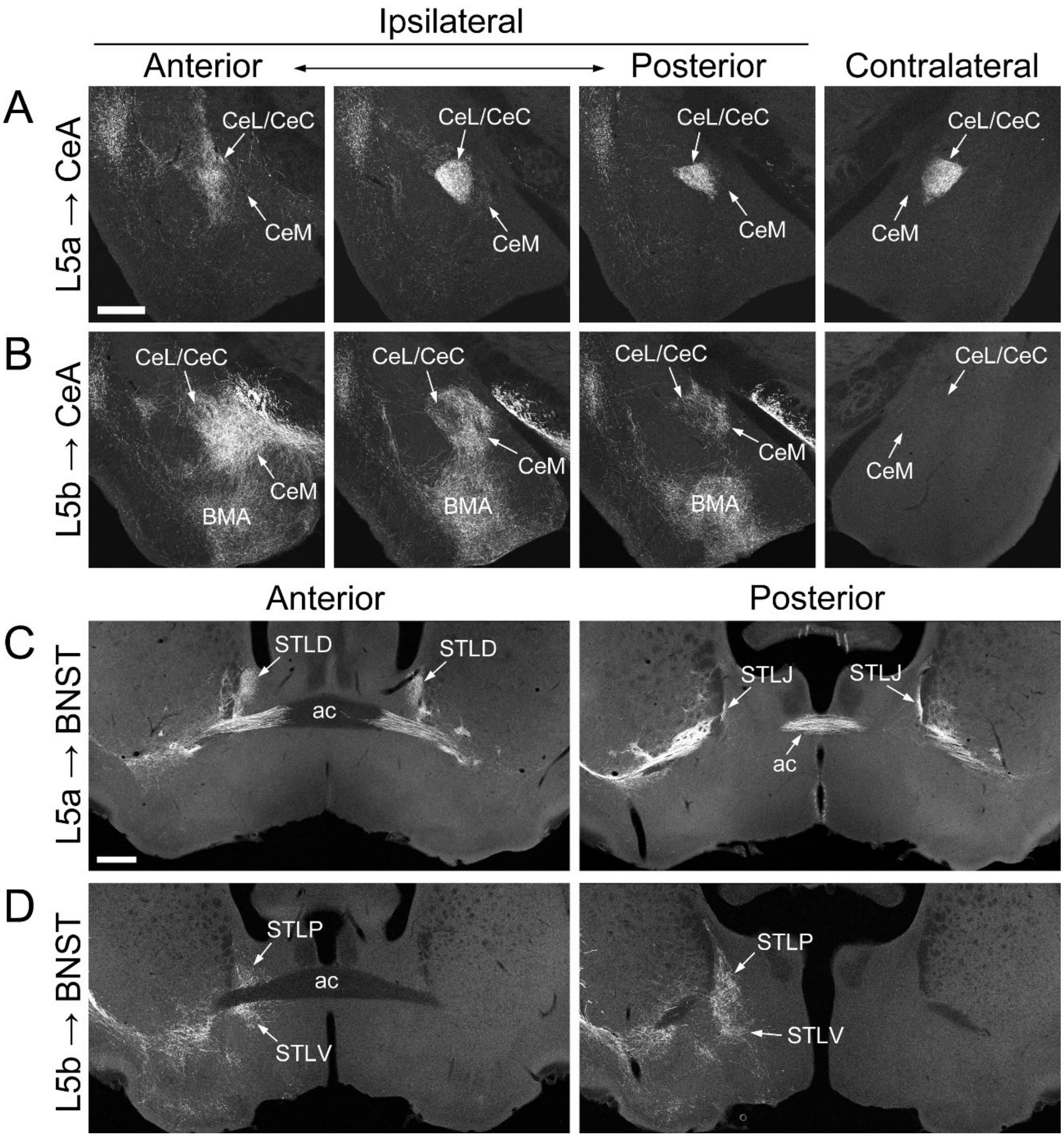


**Fig. S3. Differential organizations of axonal projections from sublaminar neuronal populations in L5 of the insula to the extended amygdala.** (A, B) GFP-expressing axons of the L5a (A) and L5b (B) neuronal populations projecting to the CeA. The ipsilateral CeA at three levels along the A-P axis and a middle part of the contralateral CeA are shown. Note the absence of L5b-derived labeled axons in the contralateral amygdala (B). BMA, basomedial amygdala. (C, D) GFP-expressing axons of the L5a (C) and L5b (D) neuronal populations projecting to the BNST. Two sections along the A-P axis are shown. Note the absence of L5b-derived labeled axons running through the anterior commissure (ac) and projecting to the contralateral BNST (D). Gray-scale images are shown. Scale bars: 0.5 mm.

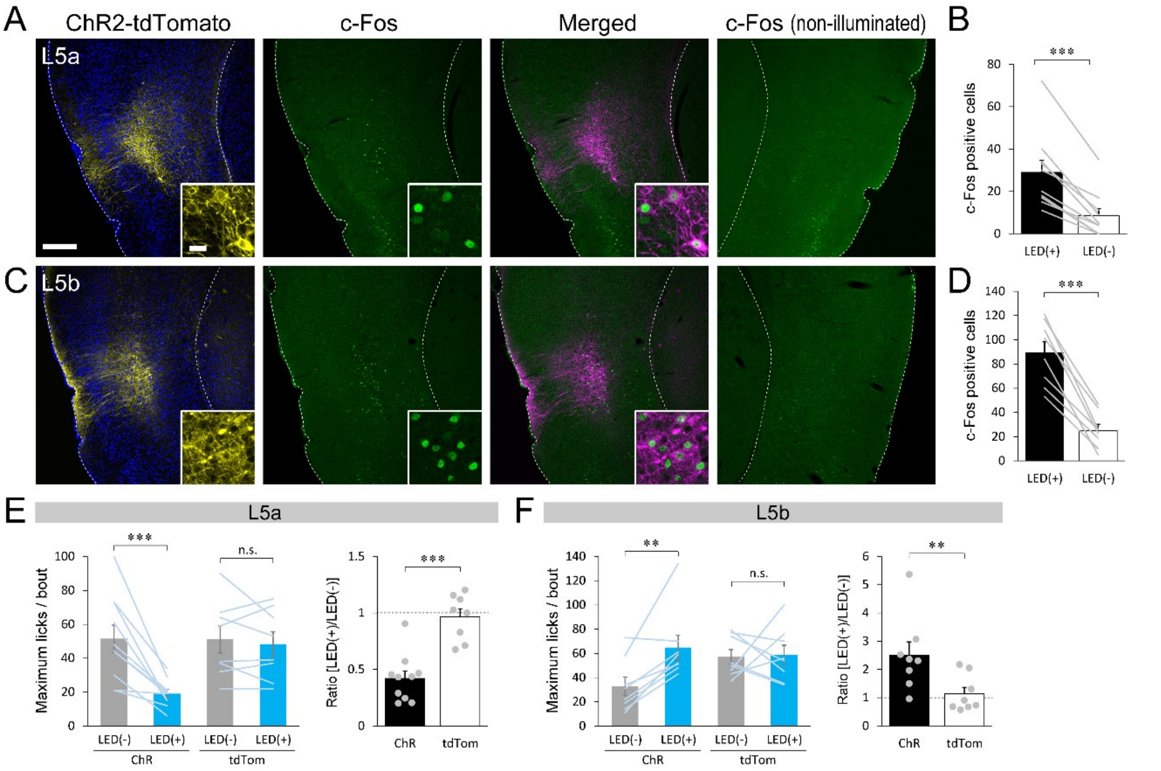


**Fig. S4. Sublayer-specific optogenetic activation of the insula and its effect on appetitive licking.** (A, C) c-Fos expression in ChR2-expressing L5 cell populations induced by LED illumination of the insula. The expression of ChR2-tdTomato fusion protein (yellow and magenta) and c-Fos (green) in L5a (A) and L5b (C) neuronal subpopulations of the left insula. Few c-Fos immunoreactive cell nuclei are seen in the right (non-illuminated) insula. Scale bars: 0.2 mm, 20 μm (inset). (B, D) Quantification of the number of c-Fos immunoreactive cells (sum of the number of cells from three sections per mouse) in the illuminated [LED(+)] vs. non-illuminated [LED(-)] side of the insula (L5a: 29.0 ± 5.7 for [LED(+)] vs. 8.5 ± 3.4 for [LED(-)], $p < 0.001$, n = 10 mice, B; L5b: 89.1 ± 9.3 for [LED(+)] vs. 25.0 ± 5.2 for [LED(-)], $p < 0.001$, n = 8 mice, D). Gray lines represent individual mice. (E, F) The maximum number of licks per bout in the single-spout test. Left: comparisons between the bouts with (blue) and without (gray) illumination for mice expressing ChR2 or tdTomato in the L5a population (ChR: 51.5 ± 8.0 for [LED(-)] vs. 19.0 ± 2.5 for [LED(+)], $p < 0.001$; tdTom: 51.1 ± 8.0 for [LED(-)] vs. 48.3 ± 7.3 for [LED(+)], p = 0.54, E) and L5b population (ChR: 32.8 ± 7.8 for [LED(-)] vs. 64.8 ± 10.3 for [LED(+)], p = 0.0055; tdTom: 57.3 ± 5.9 for [LED(-)] vs. 58.8 ± 7.9 for [LED(+)], p = 0.90, F). Right: the ratio of the maximum number of licks per bout with illumination relative to those without illumination (L5a: 0.42 ± 0.07 for ChR vs. 0.97 ± 0.07 for tdTom, $p < 0.001$, E; L5b: 2.51 ± 0.47 for ChR vs. 1.14 ± 0.23 for tdTom, p = 0.0070, F). Lines and gray dots represent individual mice (L5a: n = 10 for ChR, n = 8 for tdTom, E; L5b: n = 8 for ChR, n = 8 for tdTom, F). Dotted lines indicate the chance level. ** $p < 0.01$, *** $p < 0.001$.

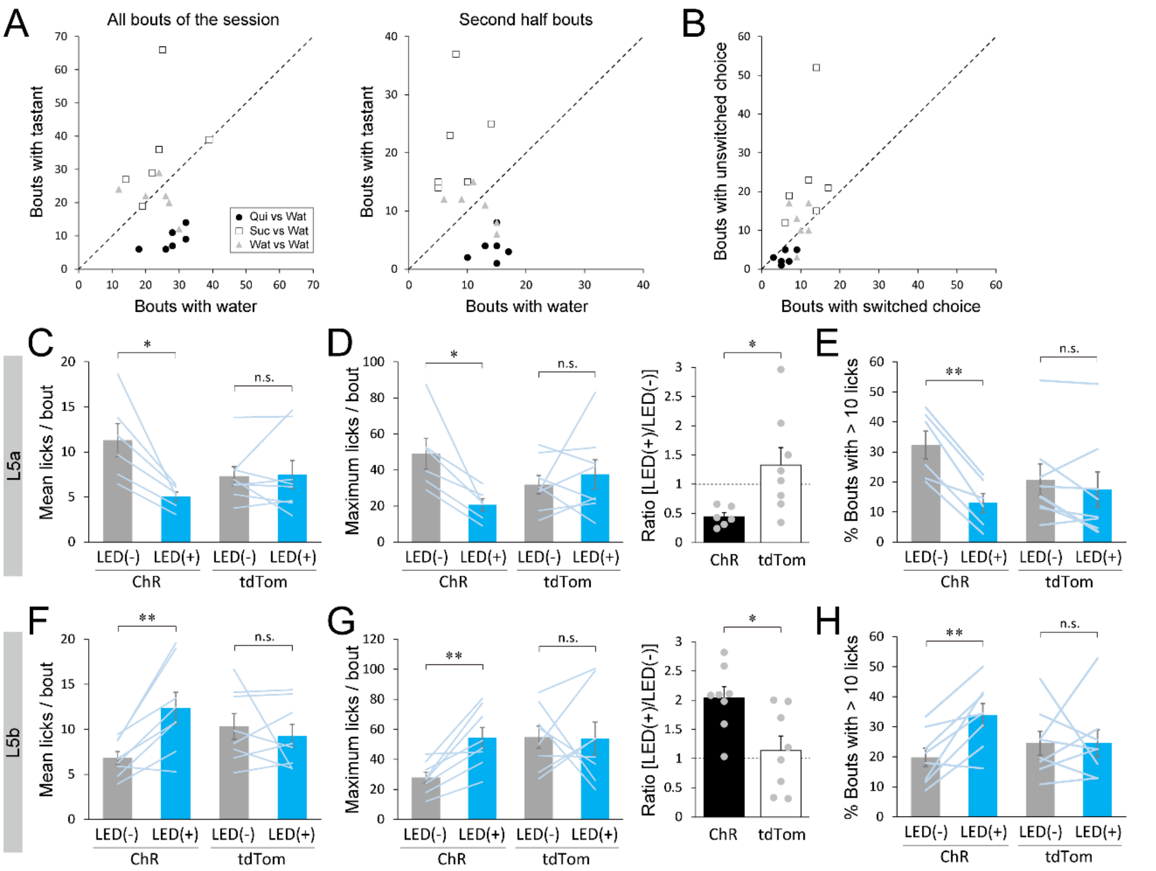


**Fig. S5. The effect of optogenetic activation of L5 subpopulations on appetitive licking in two-spout choice test.** (A) Left: the number of bouts in the whole session in which the B6 intact mouse chose the spout with water vs. tastants (quinine [Qui, black circles], sucrose [Suc, white squares], water [Wat, gray triangles]) (Qui vs Wat: 8.8 ± 1.3 for quinine vs. 27.3 ± 2.1 for water, $p < 0.001$; Suc vs Wat: 36.0 ± 6.7 for sucrose vs. 23.8 ± 3.4 for water, $p = 0.11$; Wat vs Wat: 21.5 ± 2.3 for spout-A vs. 23.2 ± 2.6 for spout-B, $p = 0.71$). Right: the same quantification within the second half bouts of the session (Qui vs Wat: 3.7 ± 1.0 for quinine vs. 14.2 ± 1.0 for water, $p < 0.001$; Suc vs Wat: 21.5 ± 3.6 for sucrose vs. 8.2 ± 1.4 for water, $p = 0.012$; Wat vs Wat: 10.7 ± 1.3 for spout-A vs. 11.5 ± 1.5 for spout-B, $p = 0.75$). N = 6 mice. (B) The number of bouts in which the mouse (B6 intact) switched vs. unswitched choice next to the bouts with tastants (Qui vs Wat: 5.8 ± 0.8 for switched spout vs. 3.0 ± 0.7 for unswitched spout, $p = 0.016$; Suc vs Wat: 11.7 ± 1.8 for switched spout vs. 23.7 ± 5.9 for unswitched spout, $p = 0.031$; Wat vs Wat: 9.8 ± 0.8 for switched spout vs. 11.7 ± 2.2 for unswitched spout, $p = 0.46$). N = 6 mice. (C, F) Comparisons of mean licks per bout between the bouts with (blue) and without (gray) illumination for mice expressing ChR2 or tdTomato in the L5a population (ChR: 11.3 ± 1.8 for [LED-] vs. 5.0 ± 0.5 for [LED+], $p = 0.010$; tdTom: 7.3 ± 1.1 for [LED-] vs. 7.5 ± 1.6 for [LED+], $p = 0.87$, C) and L5b population (ChR: 6.8 ± 0.7 for [LED-] vs. 12.4 ± 1.8 for [LED+], $p = 0.0040$; tdTom: 10.3 ± 1.4 for [LED-] vs. 9.3 ± 1.3 for [LED+], $p = 0.95$, F). (D, G) The maximum number of licks per bout in the two-spout choice test. Left: comparisons between the bouts with (blue) and without (gray) illumination for mice expressing ChR2 or tdTomato in the L5a population (ChR: 49.0 ± 8.6 for [LED-] vs. 20.7 ± 3.5 for [LED+], $p = 0.031$; tdTom: 31.9 ± 5.1 for [LED-] vs. 37.6 ± 8.1 for [LED+], $p = 0.51$, D) and L5b population (ChR: 27.9 ± 3.7 for [LED-] vs. 54.3 ± 6.9 for [LED+], $p = 0.0028$; tdTom: 54.8 ± 7.4 for [LED-] vs. 53.9 ± 10.8 for [LED+], $p = 0.95$, G). Right: the ratio of the maximum number of licks per bout with illumination relative to those without illumination (L5a: 0.44 ± 0.07 for ChR vs. 1.33 ± 0.30 for tdTom, $p = 0.021$, D; L5b: 2.04 ± 0.19 for ChR vs. 1.14 ± 0.25 for tdTom, $p = 0.013$, G). (E, H) Proportion of long drinking bouts with > 10 licks per bout in the sessions (L5a-ChR: 32.3 ± 4.6% for [LED-] vs. 13.0 ± 3.2% for [LED+], $p = 0.0012$; L5a-tdTom: 20.7 ± 5.3% for [LED-] vs. 17.5 ± 5.9% for [LED+], $p = 0.26$, E; L5b-ChR: 19.8 ± 3.1% for [LED-] vs. 33.9 ± 3.8% for [LED+], $p = 0.0037$; L5b-tdTom: 24.6 ± 4.0% for [LED-] vs. 24.5 ± 4.5% for [LED+], $p = 1.00$, H). Lines and gray dots represent individual mice (L5a: n = 6 for ChR, n = 8 for tdTom in C-E; L5b: n = 8 for ChR, n = 8 for tdTom in F-H). Dotted lines in A, B, D (right), and G (right) indicate the chance level. * $p < 0.05$, ** $p < 0.01$.

**Table S1.** List of reagents, tools, and resources used in this study.

| Reagent, tool, and resource | Source | Identifier | Concentration/dilution/titer |
|---|---|---|---|
| Drugs and reagents | | | |
| Ketamine injection 5% | Fujita Pharmaceutical Co., Ltd., Tokyo | N/A | |
| Selactar 2% Injection | Bayer Yakuhin, Tokyo | N/A | |
| Lepetan injection $0.2^{mg}$ | Otsuka Pharmaceutical Co., Ltd., Tokyo | N/A | |
| Baytril 5% Injection | Bayer Yakuhin, Tokyo | N/A | |
| Cholera Toxin Subunit B (Recombinant), Alexa Fluor™ 488 Conjugate | Thermo Fisher Scientific | C22841 | 0.5% (in 0.1 M phosphate buffer) |
| Cholera Toxin Subunit B (Recombinant), Alexa Fluor™ 555 Conjugate | Thermo Fisher Scientific | C22843 | 0.5% (in 0.1 M phosphate buffer) |
| Mowiol® 4-88 | Polysciences, Inc. | 17951 | |
| Antibodies | | | |
| Anti-c-Fos, rabbit polyclonal | Sigma-Aldrich | F7799 | 1:5000 |
| Anti-CTIP2, rat monoclonal [25B6] | Abcam | ab18465 | 1:1000 |
| Anti-FOXP2, rabbit polyclonal | Abcam | ab16046 | 1:1000 |
| Anti-FOXP2, mouse monoclonal | Atlas Antibodies | AMAb91361 | 1:2000 |
| Anti-GFP, rat monoclonal [GF090R] | Nacalai Tesque Inc., Kyoto Japan | 04404-84 | 1:2000 |
| Anti-NECAB1, rabbit polyclonal | Atlas Antibodies | HPA023629 | 1:2000 |
| Anti-PKCδ, mouse monoclonal | BD Biosciences | 610398 | 1:500 |
| Anti-RGS14, mouse monoclonal [NeuroMab clone N133/21] | NeuroMab, Davis CA | 75-170 | 1:500 |
| Donkey anti-Rat IgG (H+L) Highly Cross-Adsorbed Secondary Antibody, Alexa Fluor 488 | Thermo Fisher Scientific | A-21208 | 1:1000 |
| Donkey Anti-Mouse IgG H&L (Alexa Fluor® 647) preadsorbed | Abcam | ab150111 | 1:1000 |
| Donkey Anti-Rabbit IgG H&L (Alexa Fluor® 647) preadsorbed | Abcam | ab150063 | 1:1000 (for FOXP2) |
| Donkey anti-Rabbit IgG (H+L) Highly Cross-Adsorbed Secondary Antibody, Alexa Fluor Plus 647 | Thermo Fisher Scientific | A32795 | 1:1000 (for c-Fos) |
| Donkey anti-Rabbit IgG (H+L) Highly Cross-Adsorbed Secondary Antibody, Alexa Fluor 488 | Thermo Fisher Scientific | A-21206 | 1:1000 (for c-Fos) |
| Viruses | | | |
| rAAV2-CAG-GFP | UNC Vector Core (Edward Boyden) | N/A | 4.3x10^12 (qPCR titer vg/ml) |
| rAAV2-CAG-FLEX-GFP | UNC Vector Core (Edward Boyden) | N/A | 3.7x10^12 (qPCR titer vg/ml) |
| pAAV-Ef1a-mCherry-IRES-Cre (AAV Retrograde) | Addgene (Karl Deisseroth) | 55632-AAVrg | 1.3x10^13 (GC/ml) |
| AAV-CAG-hChR2-H134R-tdTomato (AAV Retrograde) | Addgene (Karel Svoboda) | 28017-AAVrg | 9.0x10^12 - 1.0x10^13 (GC/ml) |
| pAAV-CAG-tdTomato (codon diversified) (AAV Retrograde) | Addgene (Edward Boyden) | 59462-AAVrg | 1.0x10^13 (GC/ml) |
| Mouse | | | |
| C57BL/6JJmsSlc | Japan SLC, Shizuoka Japan | N/A | |
| Softwares | | | |
| ImageJ 1.52a | NIH | https://imagej.nih.gov/ij/ | |
| Arduino IDE 1.8.12 | Arduino.cc | https://www.arduino.cc/en/software | |
| R version 4.0.4 | The R Foundation | https://www.r-project.org/ | |
| Tools for optogenetics and behavior | | | |
| Chip LED (470 nm) | OptoSupply Ltd, N.T., Hong Kong | OSBL1608C1A | |
| Teleopto receiver | Bio Research Center Co., Ltd., Aichi Japan | TeleR-1-P | |
| Teleopto infrared emitter | Bio Research Center Co., Ltd., Aichi Japan | TeleEmitter | |
| Teleopto controller | Bio Research Center Co., Ltd., Aichi Japan | TeleRemocon | |
| Arduino UNO Rev3 | Arduino, Somerville, MA | A000066 | |
| Feeding needle | Natsume Seisakusho Co., Ltd., Tokyo Japan | KN-348-20G-50 | |
| Kwik-Sil™ silicone adhesive | World Precision Instruments, Inc., Sarasota FL | KWIK-SIL | |
| Vetbond | 3M, St. Paul, MN | 1469SB | |
| Super-Bond | Sun Medical Co., Ltd., Shiga, Japan | Super-Bond C&B | |

**Dataset S1 (separate file).** List of datasets and the statistical analyses in main and supplementary figures.
